# Dynamic gain decomposition reveals functional effects of dendrites, ion channels and input statistics in population coding

**DOI:** 10.1101/2022.02.04.479104

**Authors:** Chenfei Zhang, Omer Revah, Fred Wolf, Andreas Neef

## Abstract

Modern, high-density neuronal recordings reveal at ever higher precision how information is represented by neural populations. Still, we lack the tools to understand these processes bottom-up, emerging from the biophysical properties of neurons, synapses, and network structure. The concept of the dynamic gain function, a spectrally resolved approximation of a population’s coding capability, has the potential to link cell-level properties to network-level performance. However, the concept is not only useful but also very complex because the dynamic gain’s shape is co-determined by axonal and somatodendritic parameters and the population’s operating regime. Previously, this complexity precluded an understanding of any individual parameter’s impact. Here, we decomposed the dynamic gain function into three components corresponding to separate signal transformations. This allowed attribution of network-level encoding features to specific cell-level parameters. Applying the method to data from real neurons and biophysically plausible models, we found: 1. The encoding bandwidth of real neurons, approximately 400 Hz, is constrained by the voltage dependence of axonal currents during early action potential initiation. 2. State-of-the-art models only achieve encoding bandwidths around 100 Hz and are limited mainly by subthreshold processes instead. 3. Large dendrites and low-threshold potassium currents modulate the bandwidth by shaping the subthreshold stimulus-to-voltage transformation. Our decomposition provides physiological interpretations when the dynamic gain curve changes, for instance during spectrinopathies and neurodegeneration. By pinpointing shortcomings of current models, it also guides inference of neuron models best suited for large-scale network simulations.

**Significant Statement:** The dynamic gain function quantifies how neurons can engage in collective, network-level activity, shape brain rhythms and information encoding. Its shape results from a complex interaction between properties of different molecules (ion channels) and cell compartments (morphology, resistance), and is so far only understood for the simplest neuron models. Here we provide an interpretable analysis, decomposing the dynamic gain based on the stimulus transformation steps in individual neurons. We apply the decomposition to data from real neurons and complex models, and attribute changes of the dynamic gain to specific sub- and suprathreshold processes. Using this decomposition method, we reveal the relevance of subthreshold potassium channels for ultrafast information encoding and expose the shortcomings of even the state-of-the-art neuron models.

## Introduction

The brain’s computational abilities are realized by local networks of thousands of neurons, jointly encoding and processing information in their population activity. Understanding how this collective activity emerges from the individual cells’ properties is key to link cortical computation to molecular and cell biology. It is clear that the dynamics of action potential (AP) initiation is shaping information encoding on the network level Wolf et al. (2014), and the organelle supporting AP initiation, the axon initial segment (AIS) is a hub of plasticity with an intriguing, highly specialized molecular composition and nanostructure Huang and Rasband (2018). But a concept that connects AIS molecular structure to cellular function to network performance is lacking. Similarly, neuron gross morphology impacts encoding performance and possibly even cognitive abilities Goriounova et al. (2018), because large dendrites accelerate AP initiation and thereby enhance information encoding capacity Eyal et al. (2014); Testa-Silva et al. (2014). Theoretical neuroscience developed an abstract concept, the dynamic gain function, which connects neuronal function to population dynamics. If we can disentangle how ion channels, subcellular morphology and synaptic time scales interact to shape this dynamic gain function, it becomes a potent tool connecting molecular and cell biology to network neuroscience.

In the language of theory, the dynamic gain function is the linear response function of a population of neurons receiving a common feed-forward input (Fig. 1A1 and A2, Methods) Knight (1972). In practical terms, it captures the frequency preference of neurons, their ability to tune-in to rhythmic activity in recurrent networks Brunel and Wang (2003). Dynamic gain measurements characterize neurons in a working regime resembling *in vivo* activity, which revealed unexpected neuronal properties, such as a very wide encoding bandwidth Köndgen et al. (2008); Boucsein et al. (2009); Higgs and Spain (2009); Broicher et al. (2012); Ilin et al. (2013); Testa-Silva et al. (2014); Lazarov et al. (2018); Ostojic et al. (2015); Linaro et al. (2018); Goriounova et al. (2018); Tchumatchenko and Wolf (2011) (but see Borda Bossana et al. (2020)), and surprisingly rapid and strong flexibility Merino et al. (2021). Ultrafast, high-bandwidth encoding is likely an important determinant of the brain’s exquisite temporal performance Thorpe et al. (1996) and is suggested to underlie evolutionary pressure Lazarov et al. (2018). The rapid retuning of the frequency preference of a common interneuron class in the prefrontal cortex Merino et al. (2021) could shape thetagamma cross-frequency coupling. Dynamic gain measurements facilitate a mechanistic understanding of this phenomenon, long suspected to contribute to information routing.

**Figure 1:**
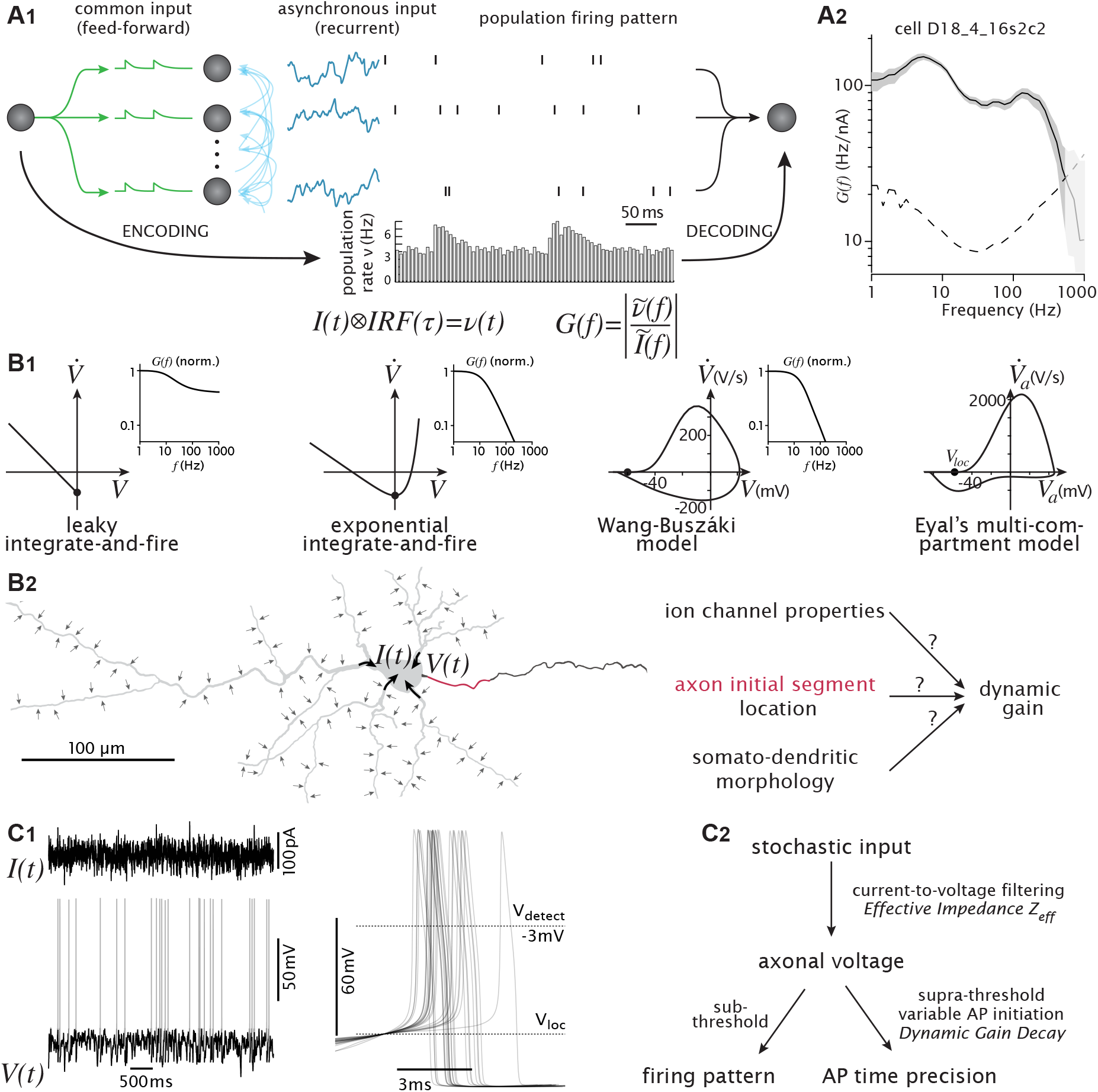
Physical signal processing steps guide dynamic gain decomposition. **A1** A population encoding scheme. Feed-forward input (green) diverges onto a recurrently connected neuronal population. The dominating, asynchronous, recurrent input causes weakly-correlated AP patterns. Nevertheless, the population rate can faithfully reflect changes in the common input. A downstream neuron can detect this signal, filtered through the population’s transfer function, the dynamic gain *G*(*f*). **A2** Experimentally determined dynamic gain function of a layer 5 pyramidal cell, obtained from 2,330 APs (data from Revah et al. (2019)). Note the wide bandwidth, and the maxima and minima. Confidence interval (grey band) and noise floor (dashed) are obtained by bootstrapping (see Materials and Methods). **B1** Evolution of neuron models’ phase plots, from point neurons without initiation dynamics to Eyal’s multi-compartment model. Locations of local minima during AP initiation are marked with dots and denoted as *V*_loc_. Normalized dynamic gain functions of the first three model variants are shown next to the phase plots. **B2** Schematic representation of neuronal signal transformation. Distributed synaptic inputs drive current into the soma and axon; the axonal voltage (*V*_*a*_) changes. For real neurons and biophysically plausible multi-compartment models, various parameters shape these transformations, attribution of dynamic gain changes to specific parameters is a challenge. **C1** Globally, Eyal’s model displays irregular firing patterns in response to stochastic stimuli. Locally, at AP onset, the initiation dynamics varies, signified by the large spread in the delay between the last positive crossing of *V*_loc_ and the AP detection voltage (0 mV). **C2** Decomposition of AP initiation process. *V*_*a*_(*t*) contains the subthreshold dynamics related to the firing pattern (time intervals of neighboring spikes when their voltages reach *V*_loc_), and the suprathreshold dynamics related to the precise, sub-millisecond AP timing (determined by the spike initiation dynamics after *V*_loc_). This decomposition can disentangle the functional effects of neuronal properties on population encoding.

Despite these clear advantages over conventional electrophysiological characterizations, dynamic gain measurements have not been widely adopted. We attribute this reluctance to two issues. First, dynamic gain measurements follow a statistical concept and require the analysis of thousands of spike times in relation to a stochastically fluctuating input. Second, there is no trivial relation between dynamic gain features and cellular properties, which are the focus of those electrophysiologists who could execute the measurements. Our study provides the tools to address these issues, to achieve the desired connection between cell- and network-level function.

Three features have already been shown to influence dynamic gain. First, the cells’ morphology Verbist et al. (2020); Zhang et al. (2022); Aspart et al. (2016); Eyal et al. (2014); Linaro et al. (2018); Brunel et al. (2001). Second, the active dynamics of AP initiation Brunel et al. (2001); Fourcaud-Trocmé et al. (2003); Naundorf et al. (2006); Wei and Wolf (2011); Huang et al. (2012); Öz et al. (2015), and third, the statistics of the neurons’ input Brunel et al. (2001); Tchumatchenko et al. (2011); Merino et al. (2021), influenced by synaptic receptor kinetics and activity correlations set by the brain state. Although the relevance of those three features for dynamic gain is established, there is no unifying theoretical approach to quantify their impact and disentangle their interactions. Here we introduce an analysis framework that guides not only the choice of working points but also allows for a detailed attribution of dynamic gain features in experimental and simulation data. Our dynamic gain decomposition follows the physical signal transformation from input currents into spike times and largely separates subthreshold processes from threshold dynamics. This enables us to interpret dynamic gain features and relate them to the underlying biophysical mechanisms, not only for simulated data, but also for recordings from cortical pyramidal neurons. We study a biophysically plausible, multi-compartment model with Hodgkin-Huxley type potassium and sodium channels, and find that the high encoding bandwidth is due to its type II excitability. A type I model counterpart fails to reproduce ultrafast population encoding. The addition of a dendrite impacts the dynamic gain primarily by shaping the impedance and only modulates the bandwidth that is determined by the excitability type. Interestingly, we find that the AP initiation dynamics limits the bandwidth in the experimental data, but not the model.

## Materials and Methods

### Integrate-and-fire models

To illustrate fundamental properties of the decomposition, we use very simple models, consisting of a single dynamical variable, the membrane voltage. In the leaky integrate-and-fire (LIF) model, the voltage changes due to an external current *I*, and an ohmic resistance of the membrane with a reversal potential *V*^*rev*^.

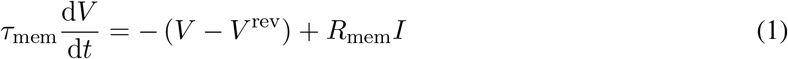

We used a reversal potential of *V* ^*rev*^ = −75 mV, and a spike threshold of −50 mV. After each spike, the voltage was reset to *V*^*rev*^. The membrane time constant *τ*_*m*_ was set to 20 ms, *R*_*mem*_ to 1.

In the exponential integrate-and-fire model voltage is additionally modified by an exponentially voltage-dependent term, representing an idealized sodium current:

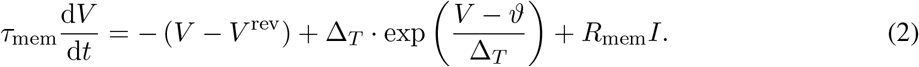

The following parameter values were used: membrane time constant and resistance *τ*_*mem*_=10 ms and *R*_*mem*_=116.417 MOhm, spike slope factor Δ_*T*_ =5 mV, threshold voltage *ϑ*=-45 mV, and reversal potential *V* ^*rev*^=-67.760304 mV. After a detection threshold of 0 mV is reached, the voltage is set to the reversal potential for 2 ms. The concrete simulations are obtained with a fifth-order Runge-Kutta-Fehlberg algorithm in Igor Pro 9 with a time step of 20 *μs*.

### Neuron morphology and biophysical properties

We used the multi-compartment model studied in Eyal et al. (2014). The neuron model is composed of one dendrite, soma and axon (Fig. 3A), and studied at 25 *μ*s time resolution. All three compartments are cylinders. Diameter and length are 20 and 30 *μ*m for soma, 1 and 50 *μ*m for the axon initial segment (AIS), and 1 and 1000 *μ*m for the myelinated axon. To examine the impact of the dendrite on population response, we studied three model variants as used in Eyal et al. (2014). The first model is without the dendrite, the second model has a dendrite with 3 *μ*m diameter and 2324 *μ*m length. The third model has a dendrite with 5 *μ*m diameter and 3000 *μ*m length of (these models are identified in Eyal et al. (2014) by axonal loads *ρ*_*axon*_ of 12, 95 and 190).

The axial resistance *R*_*a*_ is 100 Ωcm, the reversal potential for the leak current *V*_*L*_ is -70 mV, the specific membrane capacitance *c*_*m*_ is 0.75 *μ*F*/*cm^2^ and the specific leak conductance *g*_*L*_ is 3.3×10^−5^S*/*cm^2^, except for the myelinated axon, where the last two parameters are reduced to 0.02 *μ*F*/*cm^2^ and 6.6 × 10^−7^S*/*cm^2^. The dendrite, soma, and AIS, contain voltage dependent sodium and potassium channels. The sodium current is described by 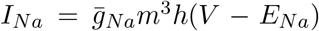 with 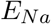 set to 50 mV. The slope factor *q*_*a*_ of the stationary gating variable *m*_*∞*_ is 9 mV. 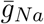 is 20 pS*/μ*m^2^ in the dendrite, 800 pS*/μ*m^2^ in the soma, and 8000 pS*/μ*m^2^ in the AIS. The potassium current is described by 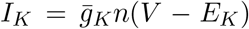 with *E*_*K*_ set to -85 mV. 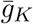 is 10 pS*/μ*m^2^ in the dendrite, 320 pS*/μ*m^2^ in the soma, and 1500 pS*/μ*m^2^ in the AIS. The dynamics of gating variables are adapted from Mainen and Sejnowski (1996) with the model temperature set to 37°C. All simulations were performed with NEURON 7.6 and 8.0 Carnevale and Hines (2006) compiled with Python 3.7 and 3.9. The code for simulations is available on github repository https://github.com/chenfeizhang/Code_Eyal_gwdg.

### Neuron model stimulation and dynamic gain function

The neuron model was driven by a current stimulus injected to the middle of the soma. To create the AP phase plots and measure the AP initiation speed, the stimulus was a constant current just above rheobase. We recorded the voltage in the AIS, 47 *μ*m away from the soma, during the first AP. For graphical comparison of the AP initiation speed of different model variants, the local minima of the phase plots were shifted to 0 mV in the x axis.

The encoding ability of the neuron population can be evaluated by injecting each neuron with an independent realization of the background noise combined with a small sinusoidal signal:

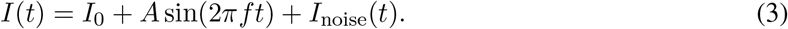

Here *I*_0_ is the mean input, *A* is the amplitude of the sinusoidal signal, and *I*_*noise*_ is a zero-mean stochastic stimulus generated by the Ornstein-Uhlenbeck (OU) process:

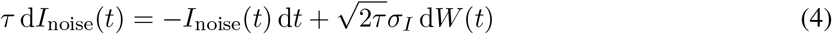

where *τ* is the correlation time, *σ*_*I*_ is the standard deviation (std), and *W* (*t*) is a Wiener process with zero mean and unit variance. The population firing rate can be expanded as:

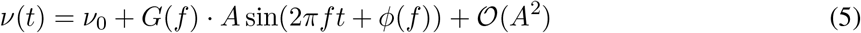

where *ν*_0_ is the mean firing rate, *G*(*f* ) is the tuning ratio between the output population firing rate and the sinusoidal signal at frequency *f, ϕ*(*f* ) is the phase shift dependent on the frequency, and *𝒪* (*A*^2^) is the higher order term of the output. We name the linear response part of the population firing *G*(·) as the dynamic gain function. The dynamic gain could be theoretically calculated in a straight forward way as the ratio of the Fourier transform of the spike output, and the Fourier transform of the current input.

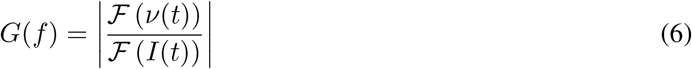

The firing rate can be understood as a sum of delta functions, located at the AP times *t*_*i*_:

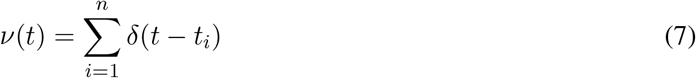

Here, we used a variant of this method, where the fraction in (6) is expanded by the Fourier transform of the input and the convolution theorem is applied. This method was proposed by Higgs and Spain (2009):

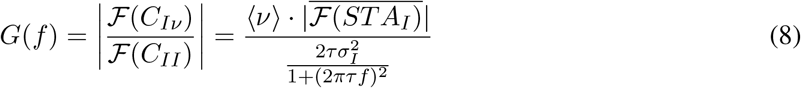

*ℱ* (*C*_*II*_) is the Fourier transform of the input auto-correlation function and *ℱ* (*C*_*Iν*_) is the Fourier transform of input-output cross-correlation. The latter is the product of mean firing rate ⟨*ν*⟩ and complex conjugate of the spike-triggered average current’s Fourier transform 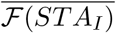. To calculate the STA input, AP times were defined as the time points at which the detection voltage was crossed from below. The default detection voltage was chosen as the voltage at which the AP waveform reaches its maximal slope. For models without such a maximum, e.g. the exponential integrate-and-fire model, the default detection threshold is chosen as a voltage, where the voltage slope is much larger than the current-induced voltage fluctuation rate.

Because the method in equation (8) does not require a sinusoidal stimulus component, we use as stimulus only the OU process defined above. Its power spectral density is known to be 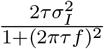, and due to the Wiener-Khinchin theorem, it equals *ℱ*(*C*_*II*_).

For each dynamic gain function, we generated 400 trials of 1000 seconds, each resulting in a 1s long STA_*I*_ input based on approximately 5000 APs. These 400 trial STA_*I*_s are averaged, yielding the final STA_*I*_ for calculating the dynamic gain function. To de-noise the dynamic gain in the high-frequency region, we applied a collection of Gaussian filters with frequency-dependent width to 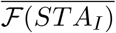, following Higgs and Spain (2009). The dynamic gain’s confidence interval was obtained as the central 95% of bootstrap dynamic gains from re-sampling the 400 STA_*I*_s 1000 times. In our graphs, it is often more narrow than the line widths. To determine the significance threshold of the gain curves, a noise floor was calculated as the 95th percentile of dynamic gain functions obtained from random AP times, obtained by shifting the original times by a random interval, larger than 1 second. We only show results above that noise floor. For a simple characterization of the shape of dynamic gain functions, we define a cutoff frequency as the point where the dynamic gain drops below 70% of the dynamic gain value at 1 Hz.

### Dynamic gain decomposition

The effective impedance *Z*_eff_(*f* ) describes, in a spectrally resolved manner, the transformation between the fluctuating current injected into the soma and the fluctuating voltage measured at the AP initiation site. Just like the dynamic gain, the effective impedance can be calculated for all frequencies at once as

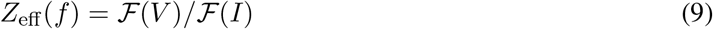

or for individual frequencies by adding a sinusoidal signal to the stochastic stimulus. The two methods yield virtually indistinguishable results. Here, we used the sine method for all multi-compartment simulations. The amplitude of the sinusoidal signal was one-fifth of the std of the stimulus. For each sinusoidal frequency, we generated 20,000 seconds of axonal voltage traces.

Whichever method is used, the challenge is to reduce the impact of spike triggered currents on the effective impedance. Two strategies can be employed, the current can be modified before the applying (9), or the voltage can be modified. In the case of simple models with reset to the reversal potential, it is very effective to add a negative current exactly at the reset time, so that the reset jump is fully explained by the modified current. If a dead-time is implemented in the model, the current after the reset is set to zero for the duration of the dead-time. In the case of more complex models with intrinsic reset mechanisms, we instead modified the voltage before applying (9). To remove the obviously non-linear AP waveforms, voltages above -50 mV were set to -50 mV. When *σ*_*I*_ is relatively small, i.e. when the neuron is operating close to mean-driven, it can happen that the spike-induced voltage excursions of after-hyperpolarization and after-depolarization exceed the input-driven voltage fluctuations. In this case, voltage excursions towards the negative values, e.g. voltages below -70 mV were also replaced by a threshold value. We averaged 20,000 pieces of 1 s of axonal voltage traces, to obtain the sinusoidally modulated average voltage. The effective impedance at the sine frequency is the ratio of the sinusoidal voltage modulation amplitude to the sinusoidal current modulation amplitude. We denote the effective impedance as *Z*_eff_(*f* ). To analyze experimental data, voltage excursions above a data-specific threshold are clipped before applying (9).

We can use the effective impedance *Z*_eff_(*f* ) to isolate the part of the dynamic gain function, which is not explained by the current-to-voltage transformation. To this end, the dynamic gain is divided by the effective impedance. The result describes the encoding of axonal voltage fluctuations into spiking in a spectrally-resolved manner, denoted as spike gain *G*_sp_(*f* ).

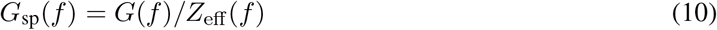

The dynamics of AP generation can vary considerably from instance to instance (Figure 1C). To explore how this variability contributes to the dynamic gain function, we change the detection threshold of APs, *V*_detect_. Importantly, we consider only the last positive crossings of *V*_detect_ before the full AP depolarization, therefore, the number of APs does not change as we study the impact of threshold. Typically, we detect APs at a voltage, where the rate of voltage rise is maximal. For Eyal’s model without the dendrite, this corresponds to -3 mV. For the experimental data, it is typically between -5 mV and + 5 mV and here we chose 0 mV for the analysis of all experimental data. When we lower this *V*_detect_, we obtain modified, earlier AP times. When we calculate *G*(*f* ) with AP times obtained with a detection voltage around AP initiation threshold, we obtain what we call the zero-delay dynamic gain *G*^0^(*f* ). For Eyal’s model, we obtained the phase plot of an AP fired just above rheobase input and used the voltage of the local minimum *V*_loc_ as an approximation of the AP initiation threshold. For experimental data, the corresponding *V*_loc_ can only be estimated. The ratio *G*(*f* )*/G*^0^(*f* ) is the dynamic gain decay caused by the variable initiation dynamics. A more extensive description with a step-by-step computation algorithm can be found at github.com/Anneef/AnTools/tree/master/Dynamic Gain Code .

### Experimental data analysis

We obtained the experimental data from Revah et al. (2019) and used the dynamic gain decomposition to interpret encoding differences between individual neurons, and between the treatment group and the control group. The dynamic gain functions were calculated with the Fourier transformation method, using the spike-triggered average current. The confidence interval was obtained as described above except that re-sampling occurred over all AP times and not trial STA_*I*_s. Noise floor calculation were performed as described above. Decomposing the dynamic gain function into effective impedance, zero-delay spike gain, and gain decay, we provided an interpretation of the encoding variability of two neurons. On the group level, *G*(*f* ), *Z*_eff_, and *G*_sp_(*f* ) were calculated for all 15 neurons of the treatment group and all 9 neurons of the control group.

The grand average within groups was obtained for the frequency range in which all individual traces were significant, i.e. above the noise floor. For the statistical analysis of *G*(200*Hz*), and *G*_sp_(200*Hz*), we tested for normality using Jarque-Bera tests and then tested for equal mean using t-tests. The phase plots for experimental data were obtained from the voltage traces by binning voltage data and then averaging the corresponding voltage derivative of each voltage point across all entries of each bin. The resulting average voltage derivative is plotted against the center of the voltage bin. The code for the analysis of experimental data is available on GitHub repository https://github.com/Anneef/AnTools.

## Results

### Decomposing dynamic gain provides subcellular resolution for the analysis of dynamic population coding

Neurons that receive a common feed-forward input constitute a population that encodes this input into changes of their population firing rate Fig. 1A1. Their encoding capability can be quantified with a linear filter, i.e the dynamic gain function *G*(*f* ). The dynamic gain’s magnitude and phase capture, in a spectrally resolved manner, how modulations of the input current cause scaled (magnitude) and delayed (phase) modulations of the population’s firing rate. Because dynamic gain reflects population-level function but is also sensitive to cellular properties, it can serve as a central link between cell-level and network-level function Brunel and Wang (2003). With the following decomposition of the dynamic gain, we make this link more accessible.

The decomposition is motivated by the simplifications that underlie the lineage of widely used simple neuron models (Fig. 1B_1_). The simplest of these models, the leaky integrate-and-fire (LIF) neuron (1), abstracts all suprathreshold dynamics as infinitely fast, and all subthreshold dynamics as a low-pass filter, i.e. an impedance *Z*(*f* ) = *R*_mem_*/*(1 + *iωτ*_mem_). We can reformulate the LIF model’s dynamic gain function *G*(*f* ) as a multiplication of a current-to-voltage transformation, captured by the impedance *Z*(*f* ), and a voltage-to-firing rate transformation, the spike gain *G*_sp_(*f*). To this end, we expand (6), the definition of the dynamic gain, with the Fourier transformation of the voltage *ℱ* (*V* ):

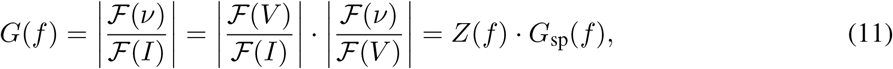

This first decomposition step is exact in LIF neurons, but how does it generalize to models with instrinsic AP initiation dynamics?

The exponential integrate-and-fire (EIF) model adds an exponentially increasing current term (2). In the subthreshold region, the voltage to current transformation becomes voltage dependent. The negative slope of the phase plot flattens, until the local minimum at *V*_*loc*_ (Fig. **2A**). Following the analysis as described in the methods, we derived the average current-to-voltage transformation of the EIF model. We found that, in the limit of low firing rates, this effective impedance converges to the impedance of a LIF model. Specifically, under stimulation conditions for which the EIF model’s voltage attains an average value of ⟨*V*⟩, the EIF model’s effective impedance converges to the impedance of a LIF model with an effective resistance *R′* = *R /* (exp ((*ϑ* − ⟨*V*⟩) */*Δ_*T*_ ) − 1) in the limit of low firing rate (Fig. 2C, dark blue).

**Figure 2:**
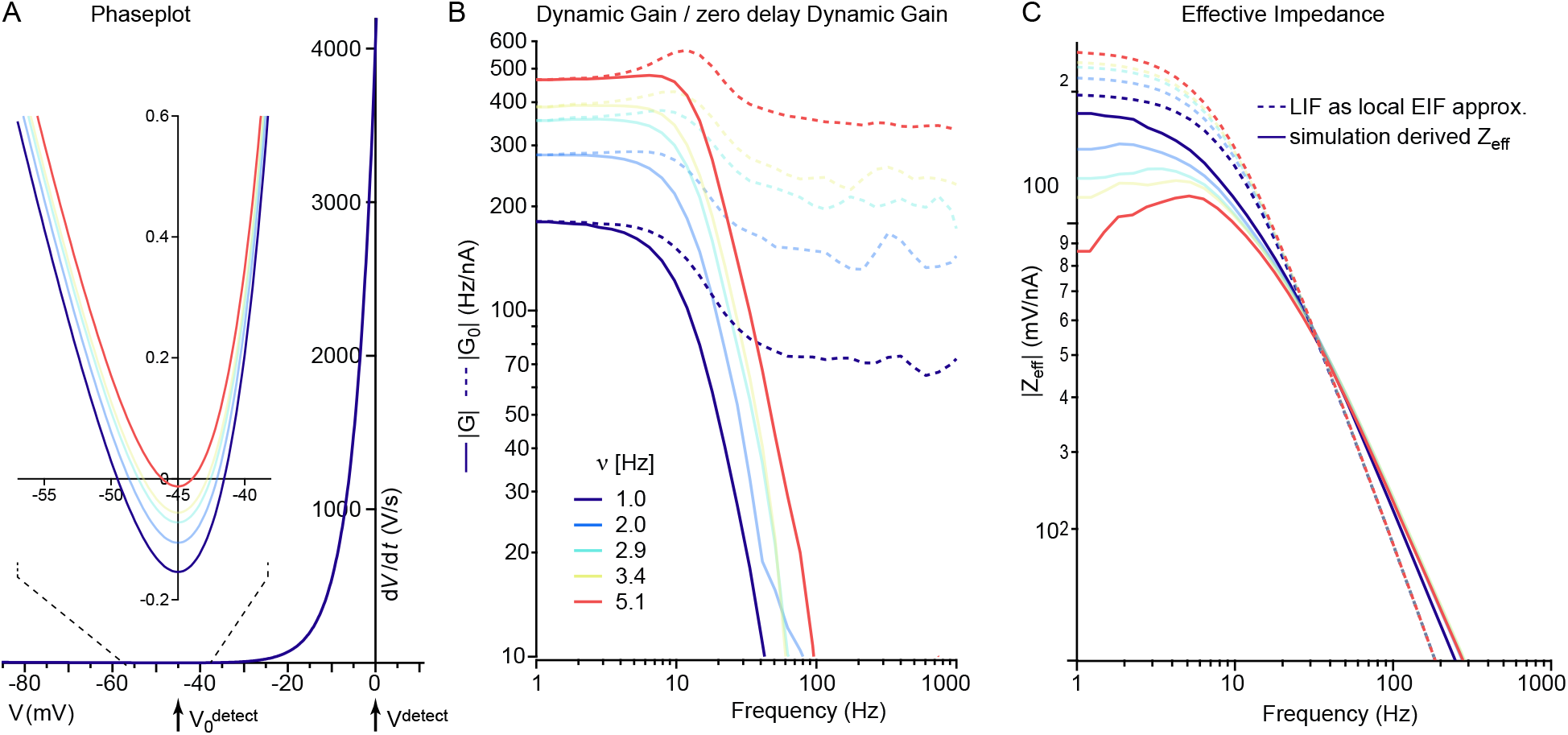
Dynamic gain decomposition of an EIF model retrieves basic building blocks corresponding to simpler LIF model variants. **A** Five phaseplots that describes the dynamics of equation (2) of the EIF model for four different current levels (I in pA: 139.3 (dark blue), 143.4, 146.3, 147.7, and 151.5 (red)). The inset magnifies the region around the local minimum. The arrows indicate the AP detection voltage used for the dynamic gain (0 mV) and the zero delay dynamic gain (-45 mV). **B** When driven with fluctuating currents with averages as in A, a standard deviation of *σ*_*I*_=15 pA and a correlation time *τ*_*corr*_=25 ms, the EIF model yields the dynamic gain displayed here in continuous lines. The corresponding zero-delay dynamic gain curves (see Materials and Methods) are show with dashed lines. The number of APs varies between the simulations (720,000 APs for 5 Hz (red), 120,000 APs for 1 Hz (dark blue), and 50,000 APs for the others, shown in fainter shades). Note how the steep decay of the dynamic gain curves turns to a flat high-frequency behavior for the zero delay dynamic gain. **C** The effective impedances for the simulations from B are displayed with continuous lines (calculated as described in Materials and Methods). The average voltage during these simulations were, in mV, -49.45, -48.86, -48.54, -48.41, and -48.17. The linear approximation of equation (2), shown by the phaseplots in A, at these voltages are the first order approximation of the current I to voltage V transformation. As described in the first section of the results, these linear approximations correspond to LIF models with modified values for membrane time constant and resistance. The impedances of these LIF models are shown with dashed lines in the color corresponding to the simulation that is approximated. Towards smaller and smaller firing rates, the numerically determined effective impedances tend toward the corresponding analytical impedances of the linear approximations around the average voltage.

Unlike the instantaneous AP “firing” of the LIF model, the more realistic AP initiation of the EIF model adds a supra-threshold component to the dynamic gain, which we can also extract. Beyond the local minimum, the phase plot’s slope d*V/*d*t* increases approximately exponentially with voltage (Fig. 2A). APs are now registered when the voltage reaches an AP detection threshold, with is much more depolarized than *V*_loc_. Between passing *V*_loc_ and reaching the AP detection threshold *V*_detect_, the dynamics is determined by a mixture of the intrinsic AP initiation current and the extrinsic stochastic stimulus, leading to variable initiation delays (see example in Fig. 1C_1_). The addition of the intrinsic dynamics causes this variable AP onset, which drastically changes the high-frequency limit of *G*(*f* ) from a constant value for LIF (Fig. 5 F) to a power law decay for EIF (Fig. 2B) Brunel et al. (2001); Fourcaud-Trocmé et al. (2003).

The simple structure of the EIF model allowed Fourcaud-Trocmé and colleagues Fourcaud-Trocmé et al. (2003) to relate the voltage dependence of the initiation current to the bandwidth of *G*(*f* ). Here we propose to study this relation with a simple, phenomenological approach that is readily generalized to models of arbitrary complexity. When we lower *V*_detect_ towards *V*_loc_, the influence of the suprathreshold dynamics is minimized and we converge to the behavior of a LIF-like model with a hard threshold and a flat gain curve.

Therefore, the gain decay of the EIF model is closely associated with the suprathreshold dynamics after *V*_loc_. In the general case, we obtain this gain decay as the ratio of two dynamic gain curves, 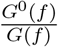, where the ‘zero-delay’ dynamic gain *G*^0^(*f* ) is obtained with ‘zero-delay’ APs detected already at *V*_loc_, while *G*(*f*) is obtained with the conventional, much more depolarized AP detection threshold (Fig. 2B). In other words, we relate the encoding capability of the full model to that of the corresponding LIF-like model version and capture the difference in the spectrally-resolved gain decay.

The introduction of other ion channel types marks the next level of model complexity relating to impedance and spike gain, represented by conductance-based models, such as the Wang-Buszáki model Xiao-Jing Wang and György Buzsáki (1996). These additional dynamical variables enable AP repolarization and richer neuronal dynamics, introducing different dynamical bifurcations at the AP threshold. This allows for a variety of firing patterns and consequently for different spike gain shapes. In spatially extended multi-compartment models (e.g. Eyal’s model Eyal et al. (2014)), currents and voltage gradients between compartments increase the complexity even further. The presence of various ion channels together with the extended morphology results in a more complex current-to-voltage transformation that we describe with an effective impedance *Z*_eff_(*f* ), which also includes the stimulus filtering along the path from the somatic stimulus source to the axonal AP initiation site.

In summary, our decomposition approach establishes three *G*(*f* ) components: 1) effective impedance *Z*_eff_ (*f* ), 2) zero-delay spike gain 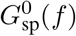 and 3) dynamic gain decay 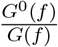. The third component captures the impact of AP initiation on the precise spike timing, reflecting the variability in the supra-threshold AP shape (Fig. 1C_1_). The first two capture subthreshold influences on *G*(*f* ), first the subthreshold transformation of current fluctuations into voltage fluctuations and then their transformation into fluctuations of the firing rate. Intrinsic frequency preferences of the neuronal dynamics can be reflected by those first two transformations, detailed in the following sections. Dynamic gain is the product of these transformations:

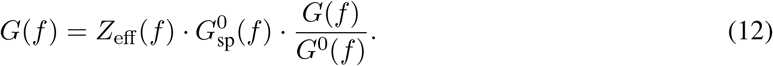

For such a decomposition, we only require the waveforms of stimulus and membrane voltage (Materials and Methods). The decomposition can therefore be performed on neuron models of arbitrary complexity and even on recordings from real neurons (Fig. 1B_2_). In the next sections, we will analyze the functional effects of biophysical and morphological parameters using experimental data and different variants of Eyal’s model, which feature a relatively wide encoding bandwidth Eyal et al. (2014). But first, Eyal’s model will be characterized and studied at different working points.

### Population encoding performance depends on the neuron’s excitability and working point

We first examined the dynamical properties of the multi-compartment model characterized by Eyal and colleagues Eyal et al. (2014), specifically the neuron model consisting of only a soma and an axon but no dendrite. In response to various constant currents injected into the soma, the firing rate *ν* of the original model displays a large discontinuity of about 32 Hz upon reaching the rheobase, indicating that it is a type II model. For comparison, we devised a model variant that shares its morphology and is equipped with the same ion channel types at the same densities and with the same voltage sensitivities. However, in the new model variant, sodium channels activate and inactivate with a 10 mV shift towards more hyperpolarized potentials (Fig. 3C). This model variant has a continuous F-I curve, indicating it is a type I model (Fig. 3B, rheobase currents aligned at 0 pA).

**Figure 3:**
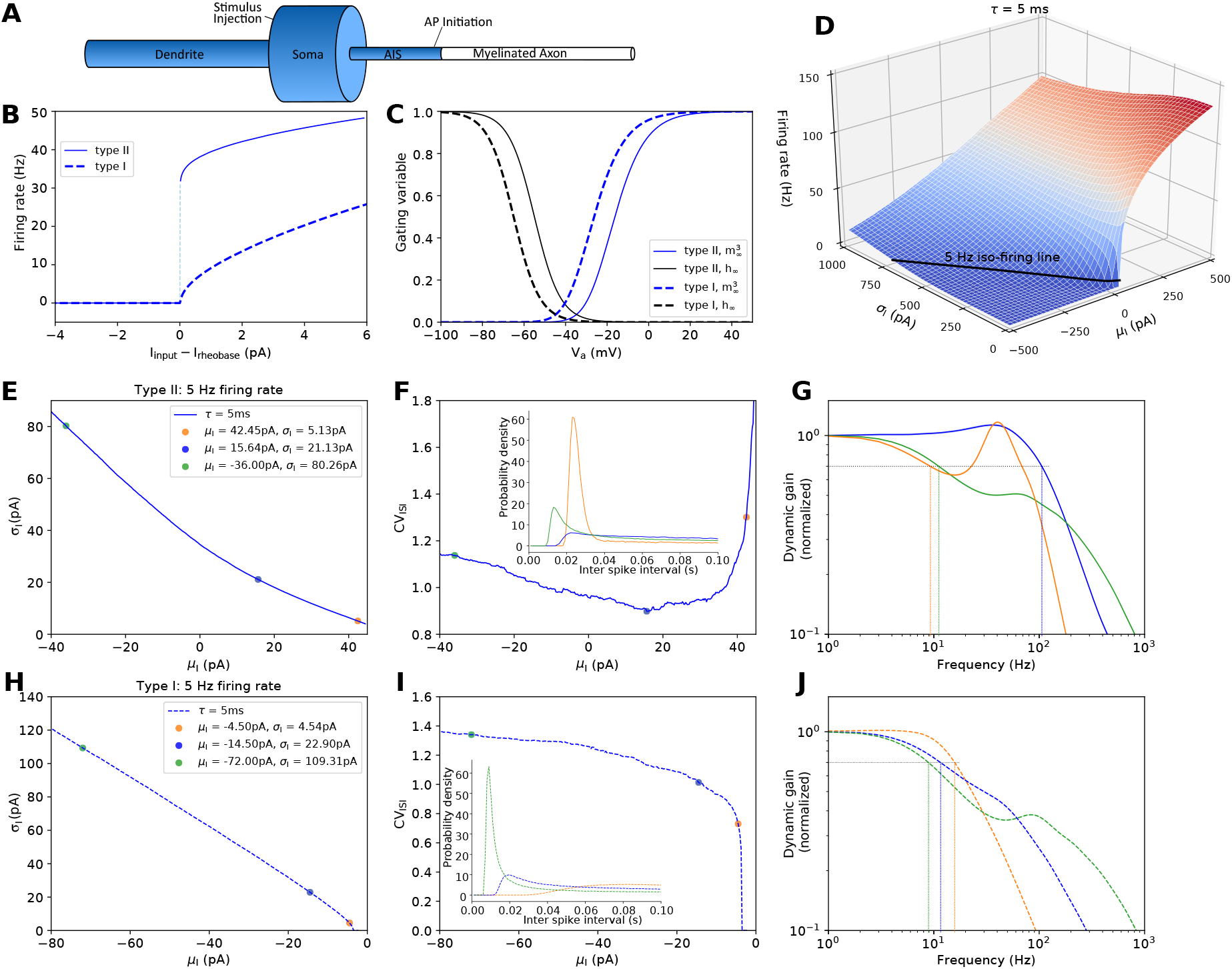
Eyal’s model can realize high-bandwidth encoding only with a type II, but not a type I excitability. **B** F-I curves with the rheobase thresholds aligned to 0 pA. **C** To switch the excitability of Eyal’s model from type II to type I, the voltage dependence of sodium channel activation and inactivation is shifted 10 mV towards more hyperpolarized potentials. *V*_*a*_ denotes the voltage at the AP initiation site. **D** 2-D firing rate surface of Eyal’s type II model as a function of mean and std of stochastic stimulus (*τ* = 5 ms). The 5 Hz iso-firing rate line is labelled black. **E** *σ*_*I*_-*μ*_*I*_ and **F** *CV*_*ISI*_-*μ*_*I*_ relation of the type II model at 5 Hz firing rate. Colored dots indicate three example working points, ranging from nearly mean-driven (orange) to fluctuation-driven (blue), and to very fluctuation-driven (green). Corresponding ISI distributions are shown in the inset panel. **G** Dynamic gain functions of the three example working points. High-bandwidth encoding is realized only when the neuron model is fluctuation-driven (blue). **H** through **J** as E through G, but for the type I model. **J** The type I model does not realize high-bandwidth encoding. Note that an increased *σ*_*I*_ enhances the dynamic gain in the high frequency region. Nevertheless, the cutoff frequency always remains below 16 Hz. For the zero-delay gains of the type I model see Extended Data Figure 3-1.

We next examined the models’ response to stochastic input with a correlation time *τ* = 5 ms. Given that the firing rate has a strong impact on the population encoding bandwidth Fourcaud-Trocmé et al. (2003), the firing rate is often fixed when different models or stimulus conditions are compared (e.g Lindner and Schimansky-Geier (2001)). Here, probing the two-dimensional firing rate surface spanned by stimulus mean *μ*_*I*_ and standard deviation *σ*_*I*_, we identified the iso-firing rate line at 5 Hz (Fig. 3D). Along this line, the neuron’s firing regime changes from nearly mean-driven to strongly fluctuation-driven. The curvature of the iso-5 Hz curve in the *σ*_*I*_-*μ*_*I*_ plane differs between the two excitability types. It is slightly concave for type I, and slightly convex for type II excitability (Fig. 3E and 3H). The firing irregularity, quantified by the coefficient of variation of inter-spike intervals *CV*_*ISI*_, differs more noticeably between the model variants. While the type I model’s firing irregularity increases monotonously with *σ*_*I*_ (Fig. 3I), the type II model displays a minimum *CV*_*ISI*_ for intermediate *σ*_*I*_ values. Towards more mean-driven conditions, *CV*_*ISI*_ increases strongly (Fig. 3F), because the intrinsic firing rate of 32 Hz is much higher than the 5 Hz target rate. In the presence of small input fluctuations around a near-rheobase average input, this low target rate is realized by bursts of near-regular rapid firing separated by long intervals. This is qualitatively very different from the irregular firing produced at working points further away from the mean-driven regime. For both model variants, we chose three working points, i.e. *μ*_*I*_-*σ*_*I*_ combinations, covering the range from mean-driven to strongly fluctuation-driven (colored dots in Fig. 3E and H). The inter-spike interval (ISI) distributions at those working points (insets in Fig. 3F and I) further illustrate the very regular firing pattern of the type II model close to the mean-driven condition. As the input fluctuations increase, the ISI distributions become more similar.

When we calculated the dynamic gain functions for the two models at the respective working points, the results were again comparable for the strongly fluctuation-driven cases, but strikingly different for other working points. When the original type II model is mean-driven (*σ*_*I*_ = 5.13 pA), the dynamic gain function appears to have a low bandwidth, dropping to 70% magnitude already at 10 Hz (orange in Fig. 3G). At intermediate and high frequencies, the shape of the dynamic gain function is dominated by a strong resonance around 40 Hz, which mirrors the peak in the ISI distribution around 25 ms. Increasing *σ*_*I*_ to 21.13 pA significantly broadens the resonance peak and increases the cutoff frequency of the dynamic gain function, which now resembles the gain curve shown in Eyal et al. (2014) (blue line). The corresponding ISI distribution is also substantially flattened, indicating the transition from mean-driven to fluctuation-driven. Interestingly, a further increase to *σ*_*I*_ = 80.26 pA, does not enhance the population encoding ability further (green line). Instead, at this strongly fluctuation-driven working point, the dynamic gain function is dominated by a lowpass filter in the low-frequency region with an apparent cutoff frequency of around 12 Hz. Towards higher frequencies, a shoulder appears and results in a local maximum around 60 Hz. The dynamic gain in the high-frequency region exceeds the other two gain curves. In parallel, the ISI distribution narrows again and moves closer to zero. The most probable ISI now is approximately 70 Hz, corresponding to the shoulder in the dynamic gain function. These results demonstrate that the encoding ability critically depends on the working point; the high encoding bandwidth reported for this type II model, is realized only in a fluctuation-driven regime with intermediate *σ*_*I*_.

The dynamic gain curves of the type I model behave strikingly different. None of the three working points lead to high-bandwidth encoding (Fig. 3J). When mean-driven (*σ*_*I*_ = 4.54 pA), the dynamic gain function has a cutoff frequency below 30 Hz, and decays with a slope of -1 in the log-log scale (orange dashed line), similar to the exponential integrate-and-fire model with standard AP initiation dynamics Fourcaud-Trocmé et al. (2003). Increasing *σ*_*I*_ to 22.90 pA to reach the fluctuation-driven regime, the dynamic gain is enhanced in the high-frequency region, while the cutoff frequency becomes smaller (blue dashed line). Further increasing *σ*_*I*_ to 109.31 pA leads to a plateau in the low-frequency region with an even lower cutoff frequency (green dash line). The dynamic gain function is larger in the high frequency with a shoulder around 100 Hz, similar to the type II model’s gain curve at the very fluctuation-driven working point (compare two green curves in F and I). Again, the ISI distribution features a peak around the time interval that corresponds to the shoulder’s frequency range. These results demonstrate that the type I model cannot reproduce high-frequency encoding throughout the biophysically plausible range of working points. In summary, we found that the encoding bandwidth can be increased tenfold, not by manipulating morphology or ion channel voltage sensitivity, but simply by increasing the potassium current around threshold to achieve a type II excitability. We next use the dynamic gain decomposition to investigate how this occurs.

### AP initiation dynamics is not the main bandwidth limitation in Eyal’s model

To decompose the models’ dynamic gains curves according to Eqn.(12), we begin with the dynamic gain decay. This component elucidates the contribution of AP initiation dynamics on dynamic gain. To prepare this analysis, we first take a closer look at the AP initiation dynamics. We previously demonstrated that high-bandwidth encoding is closely associated with the voltage sensitivity of intrinsic AP initiation dynamics, especially in the voltage region close to the local minimum of the phase plot Zhang et al. (2022). Using the voltage *V*_*a*_, recorded at the AP initiation site, we compared the phase plots of type I and type II models with their local minima (*V*_loc_) aligned at 0 mV (Fig. 4A). Although the voltage sensitivity of the sodium channel dynamics is identical for the two model variants, the voltage derivative of the type II model rises substantially more quickly out of *V*_loc_. The slower initiation of the type I model is caused by its 16 mV more hyperpolarized *V*_loc_. The sodium channels’ voltage dependence was shifted by only 10 mV, meaning that AP initiation proceeds at a voltage range with fewer activated sodium channels and of course even less potassium channels. As a consequence, the type I model’s AP initiation current displays a weaker voltage sensitivity.

**Figure 4:**
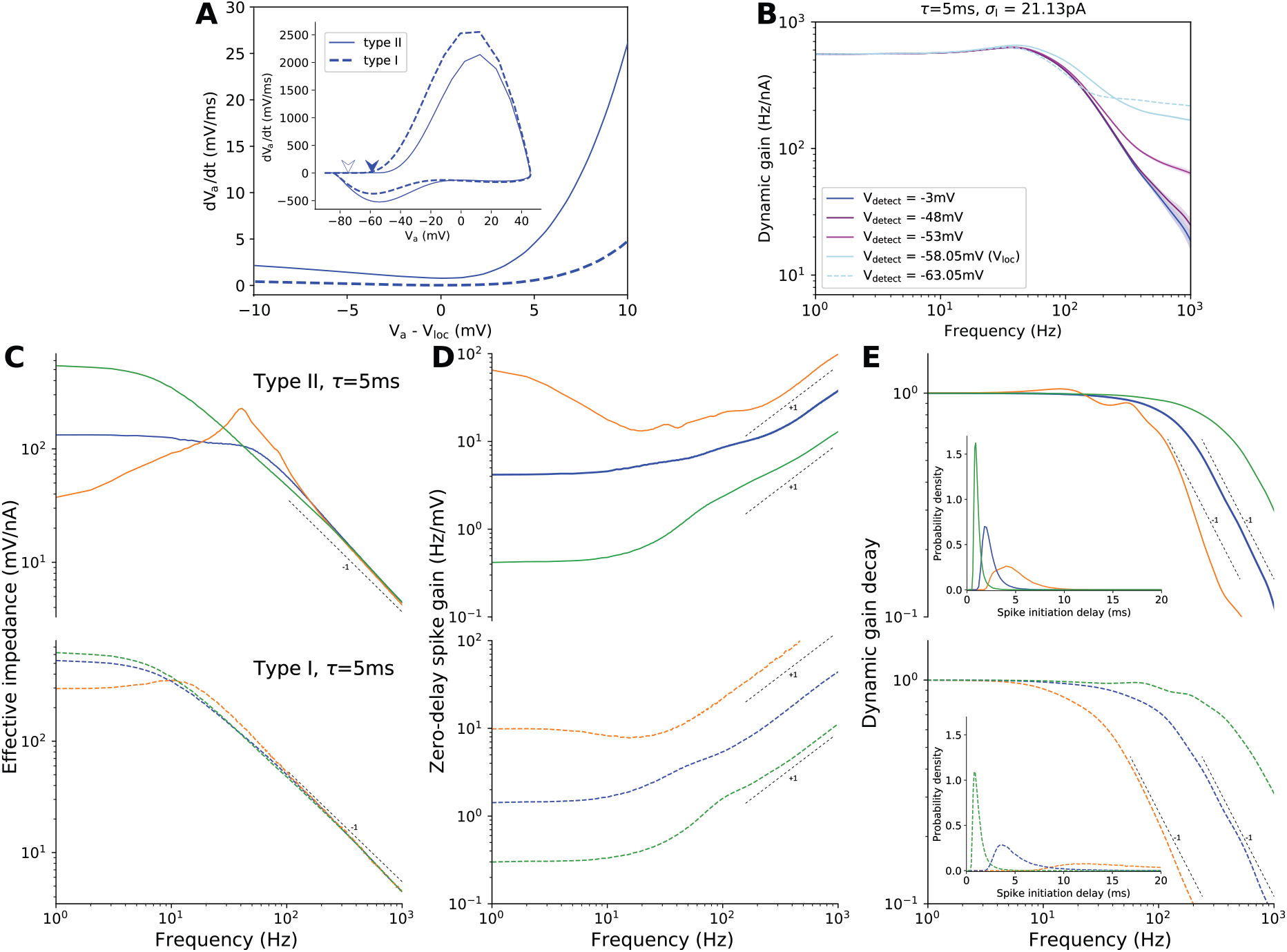
Dynamic gain decomposition based on the decomposition of the AP initiation process. **A** Insets show the AP phase plots in response to constant inputs just above rheobase. Arrows indicate the positions of the local minima *V*_loc_, of type II (-58.05 mV) and type I (-74.23 mV). The region at the beginning of AP initiation is magnified with voltages shifted to align the *V*_loc_ values to 0 mV. **B** Reducing the AP detection voltage *V*_detect_ towards *V*_loc_, enhances the dynamic gain in the high frequency region, exemplified here for the type II model. The zero-delay dynamic gain function (light blue continuous line) behaves similar to that of a LIF-like model. Further reducing the detection threshold reduces the dynamic gain in the low frequency region (light blue dash line). Note that we only include threshold crossings that continue to a full AP. **C, D, E** The dynamic gain functions from Fig. 3 F and I are decomposed into effective impedance (C), zero-delay spike gain (D) and dynamic gain decay (E). Inset panels in E are the AP initiation delay distributions at corresponding working points, color code as in Fig. 3. Effective impedance captures the transformation from somatic current to axonal voltage *V*_*a*_. Zero-delay spike gain is the ratio between zero-delay dynamic gain and effective impedance. Dynamic gain decay is the ratio between original dynamic gain and zero-delay dynamic gain (see Materials and Methods). AP initiation delay is the time interval between voltage crossing *V*_detect_ (original AP time) and the last previous positive crossing of *V*_loc_.

When we gradually lowered the AP detection voltage *V*_detect_ down to *V*_loc_, we obtained new, earlier AP times, but the number of APs stayed constant because we considered only those last positive crossings of *V*_detect_ that lead to fully developed APs. The resulting neuron model is similar to a LIF model with a hard threshold at *V*_loc_, and indeed the redefined AP times lead to a zero-delay dynamic gain function that is almost flat in the high-frequency region as expected for a LIF model (continuous light blue curve in Fig. 4B). As we detect the APs earlier, we limit the voltage range, across which the initiation dynamics and extrinsic stimulus can impact the precise AP time. Comparing the dynamic gain functions for two detection thresholds, we reveal how the interplay between intrinsic and extrinsic currents in that interval 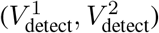 shape the dynamic gain. We find that lowering *V*_detect_ all the way from -3 mV down to -48 mV has very limited impact on the dynamic gain function. It only slightly enhances the encoding at large frequencies (above 500 Hz). However, a further decrease of *V*_detect_ by 5 to 10 mV drastically improves high-frequency encoding and has sizeable effects at frequencies down to 50 Hz, but not below. These results demonstrate that the AP initiation dynamics impacts the decay of the dynamic gain function mostly in the limit of high frequencies. It does not, however, dominate itsdecay between 40 Hz and 200 Hz, the region where it falls from its peak to its cut-off frequency. In summary, for Eyal’s model the encoding bandwidth is governed by subthreshold factors, while dynamic gain at very high frequencies is determined by the AP initiation dynamics. Consequently, the AP initiation dynamics is not the main determinant of the encoding bandwidth.

Further decreasing the detection threshold cuts into the subthreshold dynamics. The resulting changes in AP times can be substantial and affect the firing pattern and thereby also the dynamic gain in lower frequency regions (light blue dashed line in B). We can conclude that the type II model’s near constant encoding capability between 5 and 50 Hz, does not originate from its fast AP initiation after *V*_loc_. Instead, it is determined by the subthreshold dynamics before *V*_loc_.

Even the type I model’s slower AP initiation dynamics does not limit encoding between 5 and 50 Hz frequency for the fluctuation-driven regimes (blue and green). This is evident from the results of the gain decay analysis applied to the type I model (Extended Data Fig. 3-1). It is also reflected in Fig. 4E, indicating that only in the nearly mean-driven regime (orange), does the bandwidth of the gain decay limit the overall dynamic gain bandwidth.

To quantify the voltage trajectories’ variability during AP initiation, we determined the time required for each AP, to progress from the last positive crossing of *V*_loc_ to *V*_detect_ = -3 mV (see Materials and Methods). This AP initiation delay is a random variable. Its statistics are determined by the interplay between intrinsic AP initiation dynamics and extrinsic stimulus fluctuations. The initiation delay’s variance originates mainly from very variable dynamics very close to *V*_loc_, where the intrinsic currents are lowest and initiation proceeds slowest. For the type II model, 90 % of the total initiation delay are spent on crossing the first 10 mV after *V*_loc_, even though they represent only 18% of the voltage interval. These 10 mV also contribute 99 % of the delay’s standard deviation. Fig. 4B illustrates that these first 10 mV, from -58 to -48 mV, also account for the bulk of the dynamic gain decay associated with the AP initiation delay. We can use the dynamic gain decay 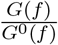 (Fig. 4E), together with the distributions of AP initiation delay, to quantify the impact of AP initiation dynamics on high-frequency encoding at the three working points in Fig. 3G and J. For the type II model, when the activity is mean-driven (orange), the AP initiation delay distribution is relatively flat with a most probable delay as large as 4 ms. Increasing *σ*_*I*_ reduces the mean of the distribution towards 0 ms (blue and green), indicating that the AP initiation delay is no longer limited by the intrinsic AP initiation dynamics. Instead, larger depolarizing inputs increase d*V/*d*t*, and accelerate the escape from the slow AP initiation region near *V*_loc_. This faster initiation is paralleled by a rightward shift in the dynamic gain decay. The bandwidth related to the initiation dynamics increases more than three-fold. For the type I model, similar transitions of the dynamic gain decays and AP initiation delay distributions can be observed when increasing *σ*_*I*_. While the two model types behave similarly when driven with very large *σ*_*I*_, towards the mean-driven working points the type I model’s AP initiation delay distribution is substantially more flattened, and consequently, the dynamic gain decay has a much lower bandwidth. We attribute this to the different firing patterns close to the mean-driven condition. The type I model produces near-regular firing without the high-frequency bursts of its type II relative. Therefore, the average AP initiation delay would approach 200 ms when *σ*_*I*_ decreases to zero and the firing rate is kept at 5 Hz. For such initiation delays exceeding the input correlation time, the input fluctuations can cause particularly large variability and consequentially deteriorate encoding precision.

### Low-frequency encoding is controlled by effective impedance and firing pattern preferences

The subthreshold part of the encoding process, which is described by the zero-delay dynamic gain *G*^0^(*f* ), can be further decomposed into effective impedance *Z*_eff_(*f* ) and zero-delay spike gain 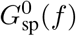 (Fig. 4C and D). The effective impedance is calculated as the Fourier transform of output voltage divided by the Fourier transform of input current (see Materials and Methods). The zero-delay spike gain is the ratio of zero-delay dynamic gain and effective impedance: 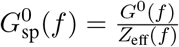.

For the type II model, the effective impedance at low frequencies varies substantially with the working point, increasing more than ten-fold at 1 Hz (Fig. 4C, upper panel). In a completely passive model without voltage-dependent conductance, the impedance does not depend on the input statistics. The observed changes in effective impedance, therefore, result from varying degrees of ion channel activity. At the mean-driven working point, the subthreshold voltage fluctuates just below *V*_loc_, where some potassium channels are already activated. Depolarization-activated potassium current opposes further depolarization, lowering the impedance. This effect is strong for low frequencies, where the potassium channel activation can follow the input changes, which explains the large drop in the orange impedance curve. At higher frequencies, the potassium channels do not actively oppose depolarization. In the limit of frequencies much higher than their activation time constant, the channels present merely a passive leak and their influence on impedance is much smaller, such that the effective impedance decays with the slope of the passive model. The impedance peak around the frequency of the intrinsic firing rate probably results from subthreshold resonance.

Increasing *σ*_*I*_ and lowering *μ*_*I*_ shifts the average voltage to regions where fewer potassium channels are activated. This increases the effective impedance at low frequency (Fig. 4C, blue continuous line) and reduces the resonance around 40 Hz. Together, these changes contribute to the 100 Hz wide bandwidth of *Z*_eff_ at the intermediate, fluctuation-driven working point. At the very fluctuation-driven working point, the sub-threshold fluctuations are even less shaped by potassium current. The effective impedance in this condition becomes a low-pass filter, determined by the passive neuronal properties (Fig. 4C, green continuous line). The type I model, in comparison, has less active ion conductances in the subthreshold range. Therefore, *Z*_eff_(*f* ) is far less sensitive to input conditions. At both fluctuation-driven working points, *Z*_eff_(*f* ) behaves similarly to a passive filter with a 10 Hz cutoff (Fig. 4C, blue and green dashed lines).

At first glance, the second subthreshold component, the zero-delay spike gain varies similarly for both models as the working points are changed. It attains larger values close to mean-driven conditions and drops with increasing *σ*_*I*_ (Fig. 4D). This overall behavior is dictated by the condition of constant firing rates. As *μ*_*I*_ decreases and *σ*_*I*_ increases, larger and larger voltage fluctuations generate the same firing rate, leading to a decrease in zero-delay spike gain. Another general feature of all zero-delay spike gain curves is their monotonic, power law increase in the high frequency limit. It compensates the decay in the effective impedance to reproduce the rather flat zero-delay dynamic gain (Fig. 4D).

Besides these general trends, the shape of the zero-delay spike gain changes between working points, in particular in the low-frequency region. For the type II model, a high plateau forms towards the mean-driven condition (orange). Conversely, a sag forms at the extremely fluctuation-driven working point (green). We will see below, that the zero-delay spike gain is the one component, that is most sensitive to the firing pattern. It represents the neuron’s propensity to turn voltage fluctuations into firing rate fluctuations based on the intrinsic dynamics. It could therefore be considered surprising, that the zero-delay spike gains of these two working points show opposite shapes. After all, both working points are characterized by bursty firing patterns, evident from the high *CV*_*ISI*_ values and the *ISI* distributions (Fig. 3F). The difference lies in the source of the bursts. At the mean-driven working point, relatively small amplitude low-frequency components drive bursts fired at the intrinsic frequency. At the very fluctuation-driven working point, large fluctuations of the extrinsic stimulus cause bursts with an intra-burst firing rate that is almost twice as high. The increased stimulus fluctuation size, together with a lower number of APs within bursts causes the zero-delay spike gain to drop at low input frequencies. This will be explored further when we analyze how repetitively fired, intra-burst APs contribute to the dynamic gain.

The decomposition of *G*(*f* ) curves across a large range of firing regimes has shown that under most conditions the initiation dynamics does not limit the overall encoding bandwidth, not even in the type I model with slower suprathreshold dynamics. The variable *G*(*f*) curves of the type II model and its ability for high-bandwidth encoding are explained by a strong stimulus dependence of *Z*_eff_(*f*) and *G*^0^ (*f*). Towards large fluctuations, both model types behave very similarly, because the extrinsic stimulus fluctuations dominate the entire signal transformation process, leading to similar *Z*_eff_(*f*), 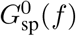 and 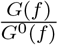. Decomposing the subthreshold and suprathreshold impact on dynamic gain, we can also aid the attribution of *G*(*f* ) changes to a particular parameter change, as we next study the influence of input correlations and neuron morphology.

### Fixing the suprathreshold impact reveals the influence of input correlations on dynamic gain

In the previous section, we studied the encoding abilities of Eyal’s type I and II models at working points ranging from mean-driven to strongly fluctuation-driven and found that only the type II model could support high-bandwidth encoding. Here we continue to examine this type II model, specifically how its encoding depends on stimulus correlation times (*τ* ). When *τ* is increased, cortical neurons, but also LIF models, show enhanced *G*(*f* ) at high frequencies, as compared to low frequencies Tchumatchenko et al. (2011); Lazarov et al. (2018); Merino et al. (2021); Brunel et al. (2001). We have called this type of input dependence the ‘Brunel effect’ and considered it an important feature of real neurons. For our study, we use the dynamic gain function in Fig. 3G (blue curve, *τ* = 5 ms) as a reference, and compare it to results obtained with more slowly fluctuating input with *τ* = 50 ms.

When we set out to define the exact working points for this juxtaposition, we concluded that *CV*_*ISI*_ is of limited practicality, because its non-monotonic *CV*_*ISI*_-*μ*_*I*_ relation in the type II model (Fig. 3F) does not allow a unique definition of working points. The dynamic gain decomposition motivates two other criteria for working point selection. While a constant *CV*_*ISI*_ aims to obtain a comparable output statistics, i.e. a comparable firing pattern, we propose to also study iso-fluctuation and iso-delay working points. They aim at a comparable subthreshold and suprathreshold dynamics, respectively. The iso-fluctuation working point is found by choosing *σ*_*I*_ and *μ*_*I*_ to fix *σ*_*V*_, the standard deviation of subthreshold voltage fluctuations. This is closely related to criteria used in several experimental studies, where *σ*_*V*_ and the target firing rate have been used to define the input parameters Tchumatchenko and Wolf (2011); Köndgen et al. (2008); Ilin et al. (2013). The iso-delay criterion controls the suprathreshold impact by fixing the average AP initiation delay. Because the intrinsic AP initiation dynamics changes not much between different working points, fixing the average initiation delay for inputs with different *τ* values, effectively fixes the impact of the extrinsic currents on AP initiation delay. Fig. 5A to B show the AP initiation delay distributions, ISI distributions and, as an inset, the *σ*_*V*_, of the working point with *τ* = 5 ms, and its iso-fluctuation and iso-delay counterparts with *τ* = 50 ms.

**Figure 5:**
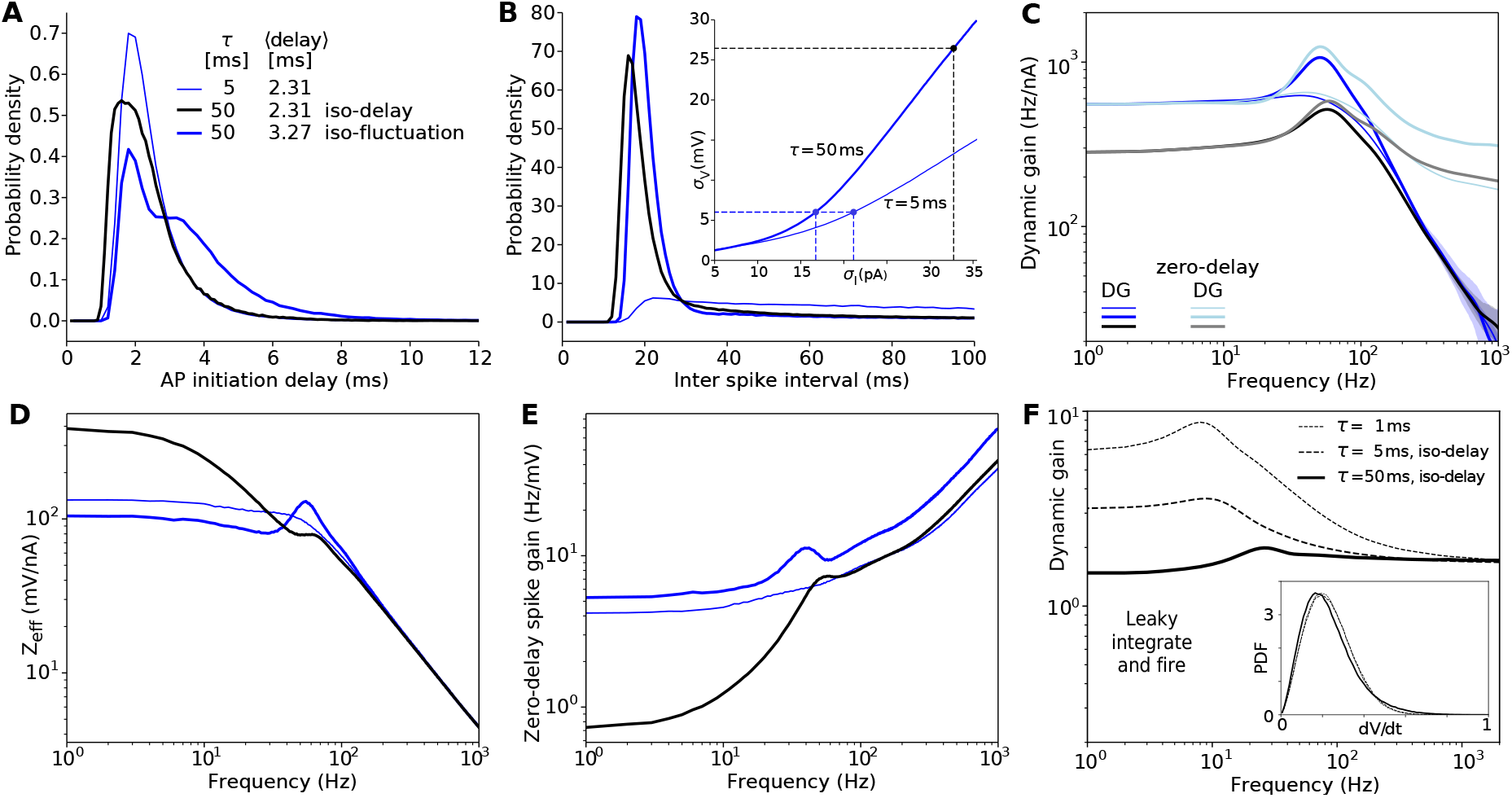
Brunel effect evaluated with two criteria fixing the sub- and suprathreshold impact on dynamic gain. **A** AP initiation delay distributions of three working points of Eyal’s type II model. Two share the same mean delay, denoted as the iso-delay working points (the thin blue curve with *τ* = 5 ms and the wide black curve with *τ* = 50 ms). In contrast, the thin and wide blue curves share the same *σ*_*V*_, denoted as the iso-fluctuation working points. This relation is indicated in the inset in B, showing the *σ*_*V*_ -*σ*_*I*_ relation for *τ* = 5 and 50 ms. The filled circles represent the three working points used from A to E. **B** to **E** ISI distributions, dynamic gain and in lighter colors the corresponding zero-delay dynamic gain functions **(C)**, effective impedances **(D), and** zero-delay spike gain functions **(E)** of the three working points above. Brunel effect, and the constituting sub- and suprathreshold contribution are different when evaluated at the iso-delay and iso-fluctuation working points. **F** For a LIF model (Materials and Methods) the iso-delay condition cannot be defined by a delay between threshold crossing and reaching a AP detection threshold, because the two coincide. However, this panel demonstrates, that an equivalent condition can be defined by matching the rate of change of the membrane voltage at the threshold crossing. If *μ*_*I*_ and *σ*_*I*_ are chosen to keep the firing rate and the average d*V/*d*t* at threshold (see inset), then the high-frequency limit of the dynamic gain does not change. This is analogous to the behavior of the zero-delay dynamic gains of the iso-delay conditions in C.

Comparing the dynamic gain functions at the iso-fluctuation working points (Fig. 5C, thin and wide blue lines), we find that increasing *τ* merely introduces a resonance around 50 Hz, consistent with the ISI distribution peak around 20 milliseconds (Fig. 5B). In the low-frequency region, the two gain curves basically coincide, suggesting that the iso-fluctuation criterion indeed equalizes the subthreshold dynamics across the two input conditions. Because the dynamic gain curves in the high-frequency region are also nearly identical, one could conclude that the Brunel effect is absent under iso-fluctuation conditions. However, a comparison of the zero-delay dynamic gain functions (Fig. 5C, wide and thin light blue curves), does show improved high-frequency encoding, for *τ* = 50 ms as compared to *τ* = 5 ms. This means that the dynamic gain decay for *τ* = 50 ms is larger, which is consistent with the more variable AP initiation delay (Fig. 5A).

Next, we examined the Brunel effect at the iso-delay working points. With the same average AP initiation delay, the overall shapes of the AP initiation distributions are also similar (Fig. 5A thin blue line and grey line). Achieving similar initiation delays with much longer input correlations requires a larger *σ*_*I*_. *σ*_*V*_ is even more than quadrupled from 6 to 26 mV (inset in Fig. 5B). But even the drastic increase in voltage fluctuations shifts the ISI distribution only slightly to smaller values (Fig. 5B wide lines). The large increase in *σ*_*I*_ for *τ* = 50 ms results in a strong reduction of the dynamic gain function in the low-frequency region (Fig. 5C). It now lies below the dynamic gain function for *τ* = 5 ms. The high-frequency limits of the two *G*(*f* ) curves are close to each other, and even the zero-delay gain functions *G*^0^(*f* ), now attain similar values in the high-frequency limit. We conclude that the type II model reproduces the Brunel effect at the iso-delay working points, which fix the impact of suprathreshold dynamics on population encoding. In this condition, the relative improvement of high-frequency encoding observed in the normalized dynamic gain functions is realized by reducing the unnormalized dynamic gain in the low-frequency region when *τ* is larger. At iso-fluctuation working points, on the other hand, the low-frequency encoding is nearly constant and no Brunel effect manifests in the dynamic gain curves because the stronger dynamic gain decay cancels increased zero-delay dynamic gains at *τ* = 50 ms.

We can now use the dynamic gain decomposition to check whether iso-fluctuation and iso-delay criteria indeed standardize sub- and suprathreshold contributions, and to further understand the source of the Brunel effect (Fig. 5D and E). We find that the effective impedances and zero-delay spike gains are very similar at the iso-fluctuation working points, in accordance with its purpose of fixing subthreshold dynamics. Within the identical subthreshold voltage fluctuation range, the active ion conductances recruited for stimulus filtering are close to each other. The remaining difference probably stems from the slightly more depolarized average voltage for *τ* = 50 ms, leading to slightly higher potassium current activation. Two zero-delay spike gain curves are almost paralleled with each other, indicating that the voltage frequency components are transformed into firing frequency components in similar ways for both correlation times. However, at the iso-delay working points, the effective impedance and spike gain in the low-frequency region show large but opposite changes in response to increasing *τ* . While the effective impedance at 1 Hz is more than 3 times larger for *τ* = 50 ms, the spike gain at 1 Hz is more than 5 times larger for *τ* = 5 ms. The difference in effective impedance is readily explained by the difference in average potassium channel conductance. *σ*_*V*_ is about 6 mV for *τ* = 5 ms, and increases to about 26 mV for *τ* = 50 ms. The substantially larger fluctuations into the hyperpolarized range cause ion channel deactivation, a lower membrane conductance and thereby a larger effective impedance. This is reflected in the much lower cutoff frequency of 8.3 Hz for *τ* = 50 ms as compared to the 56 Hz for *τ* = 5 ms. Low-pass effective impedance keeps more low-frequency components of stimulus in the voltage fluctuations. As a result, the zero-delay spike gain at the iso-delay working point is drastically reduced in the low-frequency region (Fig. 5E, black curve). The decomposition of zero-delay dynamic gain functions implies two different ways to realize high-bandwidth encoding for two correlation times of input, by influencing the firing patterns emitted by the neuron. How that impacts the dynamic gain will become clear, when we next study the differential encoding capacity of fluctuation-driven APs and repetitively firing APs.

### Decomposing dynamic gain functions into repetitive firing and individual AP components

Our previous arguments suggest that the firing pattern, in itself, is an important contributor to the shape of the dynamic gain function. It could even be thought that the type II model is capable of high-bandwidth encoding in part due to its ability to fire high-frequency repetitive APs. We next attempt to analyze this more stringently, by decomposing the dynamic gain function into two parts, contributed by two AP populations (Fig. 6A).

**Figure 6:**
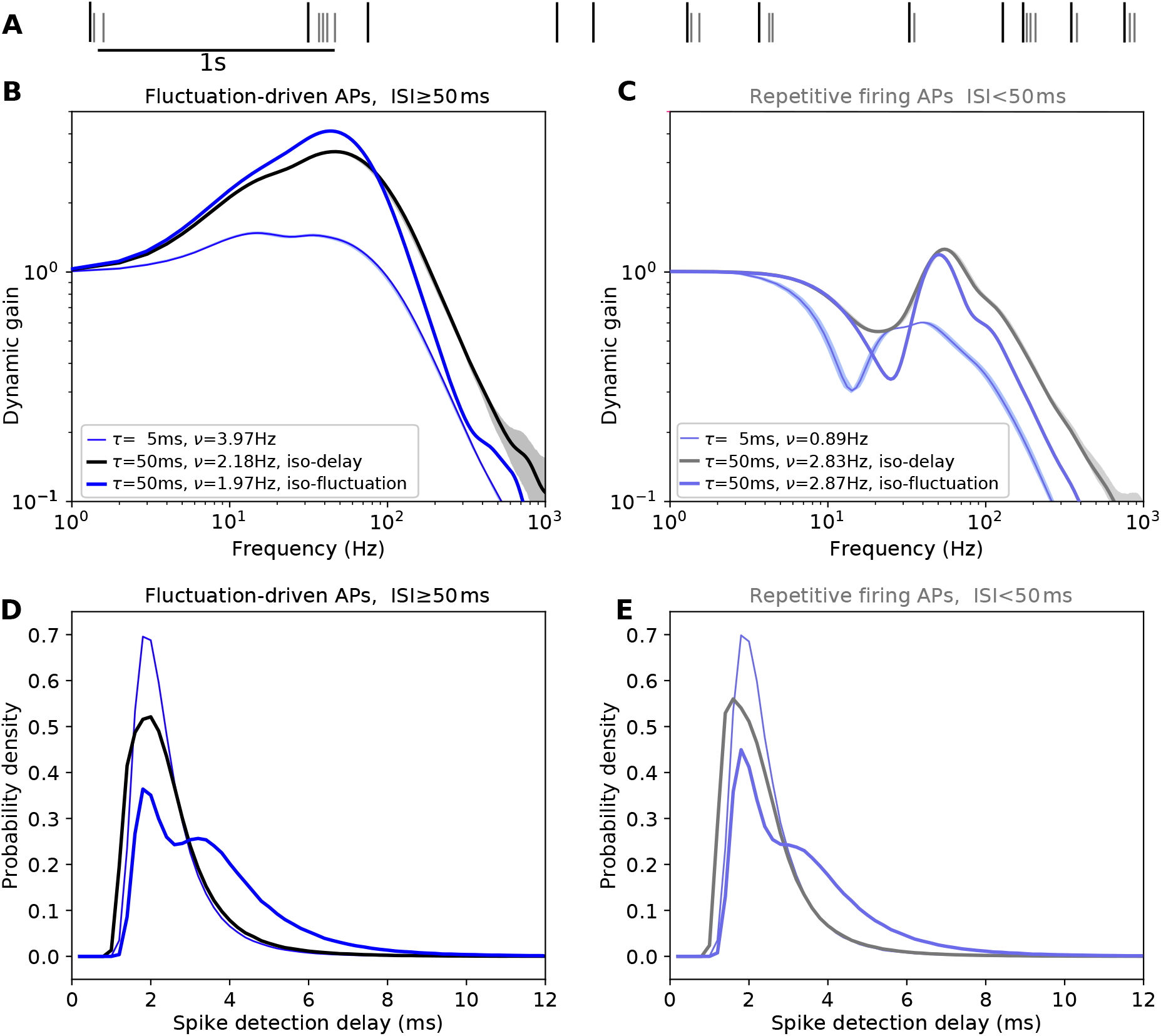
Intrinsic high-frequency firing undermines high-bandwidth encoding. **A** A short sequence of APs from the simulations in Fig. 5 (*τ* = 50 ms, iso-delay), shown in a raster plot. The APs are classified into two groups according to the preceding ISI (ISI ≥ 50 ms long, black dashes, ISI *<* 50 ms shorter, grey dashes). Separate dynamic gain functions of the iso-delay and iso-fluctuation dynamic gain functions in Fig. 5C are calculated for both spike classes and shown here. **B** The dynamic gain curves derived from fluctuation-driven APs. For the slower fluctuations, high frequency input components are encoded better than low frequency components as a consequence of resonant firing. **C** The dynamic gain curves derived from APs fired in close succession, e.g. repetitively fired APs. Resonance effects cause the dips and peaks (see main text). **D** and **E** The AP initiation delay distributions of the two spike classes are very similar in the case of iso-delay, and hence very similar to the joint distribution in Fig. 5A. The delay distributions in the iso-fluctuation case show a much longer tail. Because the typeI model produces much less bursting, the detrimental effects on encoding are weaker as shown in Extended Data Figure 6-1.

One population is fluctuation-driven APs, defined as those fired after at least 50 ms of silence. These Aps are likely to be the first one, or the only one, fired in a voltage upstroke, while the rest of APs are likely part of a burst later in an upstroke, denoted as repetitive firing APs. Fig. 6A is an illustration of the AP classification, the relative fraction of the two components is stated in Figs. 6B and 6C between correlation times. When *τ* = 50 ms, the ISI distribution is significantly more peaked around 25 ms (see Fig. 5B), and more than half of the APs are classified as repetitively firing APs. This fraction drops to 20% for *τ* = 5 ms, because bursts are terminated by the rapid *V*_*a*_ fluctuations.

We now calculated dynamic gain functions based on APs from only fluctuation-driven or only repetitive firing APs. When their phases are taken into account, these two dynamic gains can be summed to yield the curves in Fig. 5C (black and thin blue). The fluctuation-driven APs alone encode the input with a higher bandwidth, as compared to all APs together (thin blue line in Fig. 6B vs iso-delay working points in Fig. 5C). The corresponding cutoff frequencies are 133 Hz vs 105 Hz for *τ* = 5 ms, 234 Hz vs 166 Hz for *τ* = 50 ms. The shapes of these gain curves reflect the broad 20 to 50 Hz frequency encoding preference of the fluctuation-driven APs. When the intrinsic preference is overridden by very strong external fluctuations, this preference is suppressed (see the Extended Data Fig. 6-1A and C).

The repetitive firing APs alone lead to dynamic gain curves with very pronounced resonance around the preferred firing rate and a pronounced dip around half of that frequency (Fig. 6C). The dip is a direct consequence of the resonance because APs locked to an input component at the resonance frequency and fired in a short burst will be locked worse than random to half of that frequency. The dynamic gain functions derived from repetitive APs alone have a comparatively narrow bandwidth, especially for *τ* = 5 ms, where the cutoff frequency is below 8Hz. In conclusion, the two dynamic gain functions for *τ* = 5 and 50 ms at the iso-delay working points realize high-bandwidth encoding in different ways. One reduces the fraction of repetitive firing APs, while the other forms longer bursts of repetitive firing APs to weaken its low-pass filter effect. In either case, our results clearly show that the high encoding bandwidth of Eyal’s model with type II excitability is not a direct result of the burst firing. APs fired within bursts only contribute the pronounced peaks at the preferred firing rate. We can also use this AP classification strategy to further investigate the variation of spike gain shape across working points. Low frequencies of only a few Hertz were particularly well represented in the zero-delay dynamic gain curve, when the neuron was nearly mean-driven (Fig. 4D, orange). As the Extended Data Fig. 6-1 B shows, this is caused by a relatively large fraction of repetitive firing APs. They occur in long bursts and thereby limit the encoding of intermediate frequencies. This can be understood when considering the example of a 100 ms long burst, for which first and last APs fall on diametrical phases of a 5 Hz stimulus component and thereby limit encoding of this frequency. In contrast, the fluctuation-driven APs, do not show such a preference for very low frequencies, but they make up only a small fraction of all APs at the nearly mean-driven working point.

### Dendrites enhance high-bandwidth encoding by suppressing low-frequency effective impedance

As an application of the dynamic gain decomposition, we disentangled the complicated effects of dendritic morphology on each stage of information processing. This analysis elucidates the impact of dendrites on population encoding. We took the two model variants with a median and a large dendrite from Eyal et al. (2014) for comparison (see Materials and Methods), both display type II excitability (see inset Fig. 7A)). We first examined the impact of dendrite size on intrinsic AP initiation dynamics. Increasing the dendrite size increases the lateral current from the initiation site towards the soma, due to the larger somato-dendritic current sink. *V*_loc_ is shifted to slightly more depolarized values from -58.05 mV to -54.72 mV and -53.61 mV. As a result, the AP onset is shifted towards voltages at which the sodium channels have steeper voltage dependence. Consequently, at later stages, e.g. 10 mV/ms, the slope of the phase plot is actually larger for models with larger dendrites, as has been previously reported in Eyal et al. (2014) (see Fig. 7A). However, when aligning *V*_loc_ to (0 mV, 0 mV/ms), we found that a larger dendrite does not accelerate AP initiation close to *V*_loc_, instead, the local slope of the phase plot is reduced. A first evaluation of the subthreshold effects of the dendrites can be gained from the *σ*_*V*_ - *σ*_*I*_ relation (Fig. 7B, inset panel). The large current sink of the dendrite drastically decreases the impedance of the neuron, reflected by the reduced slopes. Compared to this drastic impedance effect, the dendrite’s impact on the suprathreshold dynamics appears to be subtle. Therefore, to compare the population encoding capabilities of the three models, we chose iso-delay working points to harmonize the suprathreshold contribution and focus on different subthreshold contributions of the dendrite.

**Figure 7:**
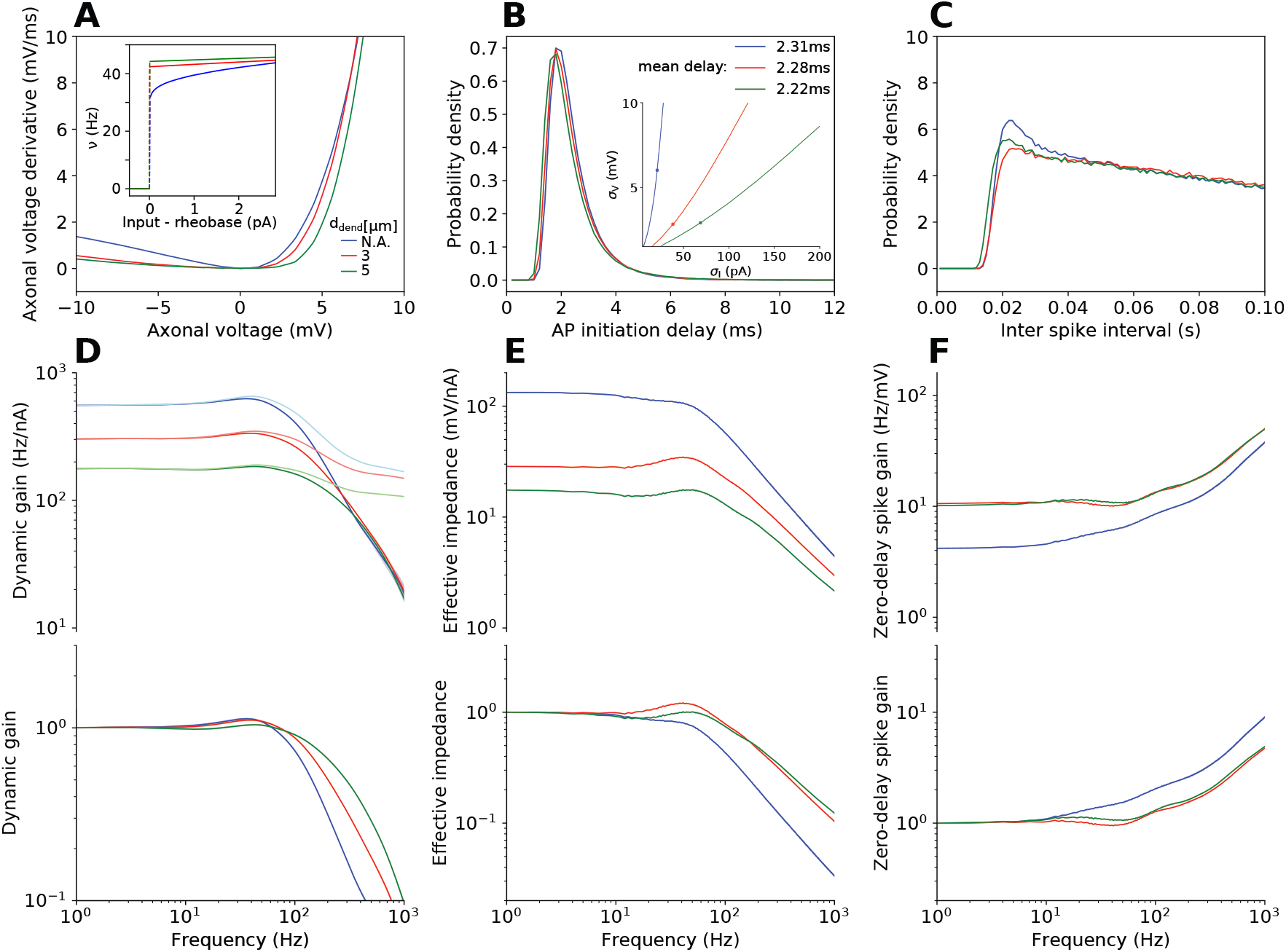
A larger dendrite shunts low-frequency inputs, but improves low-frequency spike gain. **A** Phase plots of three model variants with different dendrite sizes, *V*_loc_ aligned at (0 mV, 0 mV/ms). The inset shows the f-I curves of Eyal’s type II models with different dendrite sizes. Larger dendrites increase the minimal firing rate **B** AP initiation delay distributions at the iso-delay working points (*ν* = 5 Hz). Inset panel shows corresponding *σ*_*V*_ on *σ*_*V*_ -*σ*_*I*_ relations. **C** ISI distributions at the iso-delay working points. **D, E, F** Dynamic gain functions and **in lighter colors the corresponding** zero-delay dynamic gain functions (D), effective impedances (E) and zero-delay spike gain (F) of the three model variants calculated at iso-delay working points (upper panels), together with their normalized gain curves (lower panels). See the Extended Data Figure 7-1 for the results with *τ* = 50 ms.

Taking the dynamic gain function of *τ* = 5 ms in Fig. 5 for reference, we fixed the average AP initiation delay for the three model variants, leading to very similar distribution shapes (Fig. 7B). Under this criterion, *σ*_*V*_ is smaller for the neuron model with a dendrite (scattered dots in inset panel), and the three ISI distributions are similar to each other (Fig. 7C). Compared to the dynamic gain function without the dendrite (blue), the other two gain curves (red and green) have smaller dynamic gain below 200 Hz. In the high-frequency region, gain curves decay with similar trends, and corresponding zero-delay dynamic gain functions are also close to each other (Fig. 7D, upper panel). Normalizing the dynamic gain functions, we observed an enhancement of high-frequency encoding with a larger dendrite (lower panel in D).

Decomposing the zero-delay dynamic gain functions into effective impedances and zero-delay spike gains, we found that adding a dendrite reduces the effective impedance mostly for low frequencies, but not in the high-frequency limit. (Fig. 7E, upper panel). Low-frequency components are suppressed more strongly, because they charge a larger portion of the dendrite, while high frequencies experience much stronger spatial filtering and charge only the proximal dendrite. When the effective impedances are normalized (lower panel in E), the dendrites’ suppression of low frequencies appears as a boost of high-frequency representation in the voltage. Interestingly, a further substantial increase in the dendrite size hardly affects the shape of the effective impedance. The zero-delay spike gain curves are relatively flat in the low-frequency region, and they increase in similar trends in the high-frequency region (Fig. 7F, upper panel). The curves for the two dendrite-bearing models are almost identical (red and green), while the zero-delay gain curve of the original model has lower values (blue) because it produces very similar firing patterns (Fig. 7C) with approximately three times larger subthreshold voltage fluctuations (inset in Fig. 7B). The normalized curves in the lower panel show that, the dynamic gain enhancement caused by the effective impedance is slightly undermined at the stage of zero-delay spike gain (from blue to red, and green). Taken together, the improved high-frequency encoding in the presence of a dendrite is primarily due to a suppression of low frequencies in the effective impedance, with a minor effect of the accelerated AP initiation, leading to weaker dynamic gain decay. Studying the effect of the dendrite under slower input correlations, led to the same conclusion. In particular, the dendrite did not change the Brunel effect (Extended Data Fig. 7-1).

### Dynamic gain decomposition provides subcellular dissection of a pathophysiological insult on population coding

The dynamic gain decomposition is designed to be applicable to any neuron, independent of its complexity. It should be applicable even to recordings from real neurons, provided that the recorded somatic voltage contains sufficient information. For the subthreshold analysis, this is very likely, but the utility of somatic recordings for initiation-related analysis is not clear a priori. We therefore set out to apply the dynamic gain decomposition to experimental data and chose a specific data set from a previous study for two reasons. First, the two groups in the data set had been reported to have different input resistances, prompting us to expect different effective impedances. Second, the treatment group, but not the control group, contained neurons with very different dynamic gain curves, which we would like to understand better by decomposition.

The data in question were recorded from layer 5 pyramidal neurons in coronal slices of mouse somatosensory cortex and originally published in Revah et al. (2019). Only slices in the treatment group underwent two brief hypoxic episodes that induced spreading depolarization. Unlike simulations, experiments record voltage at the soma and not the initiation site. We showed earlier, that the transfer impedance from soma to initiation site does pose an additional filter for higher input frequencies. However, for distances up to 50 *μ*m, this effect was very small Zhang et al. (2022), and hence we speculate that using the somatic voltage will still allow decomposition.

We decomposed the dynamic gain functions of each pyramidal neuron into effective impedance and spike gain. For two, very differently affected neurons from the treatment group of Revah et al. (2019), the results are shown in Fig. 8. Neuron 1 appears very similar to control neurons, it displays encoding with a bandwidth of approximately 350 Hz (Fig. 8A). Its dynamic gain’s confidence interval widens substantially, as the gain curve drops below the noise floor. In comparison, the dynamic gain of neuron 2 (Fig. 8B) is substantially larger in the low-frequency region, decays more steeply at intermediate frequencies, and displays a lower cutoff frequency of around 200 Hz. But surprisingly, above 400 Hz, its dynamic gain reemerges above the noise floor and increases with a smaller bootstrapping confidence interval as compared to neuron 1. Also note that the absolute magnitude of the dynamic gain, for instance at 200 Hz, is higher in neuron 2, although its cutoff frequency is lower. The underlying properties that cause these differences became clear, when we decomposed the dynamic gain into the sub- and suprathreshold components.

**Figure 8:**
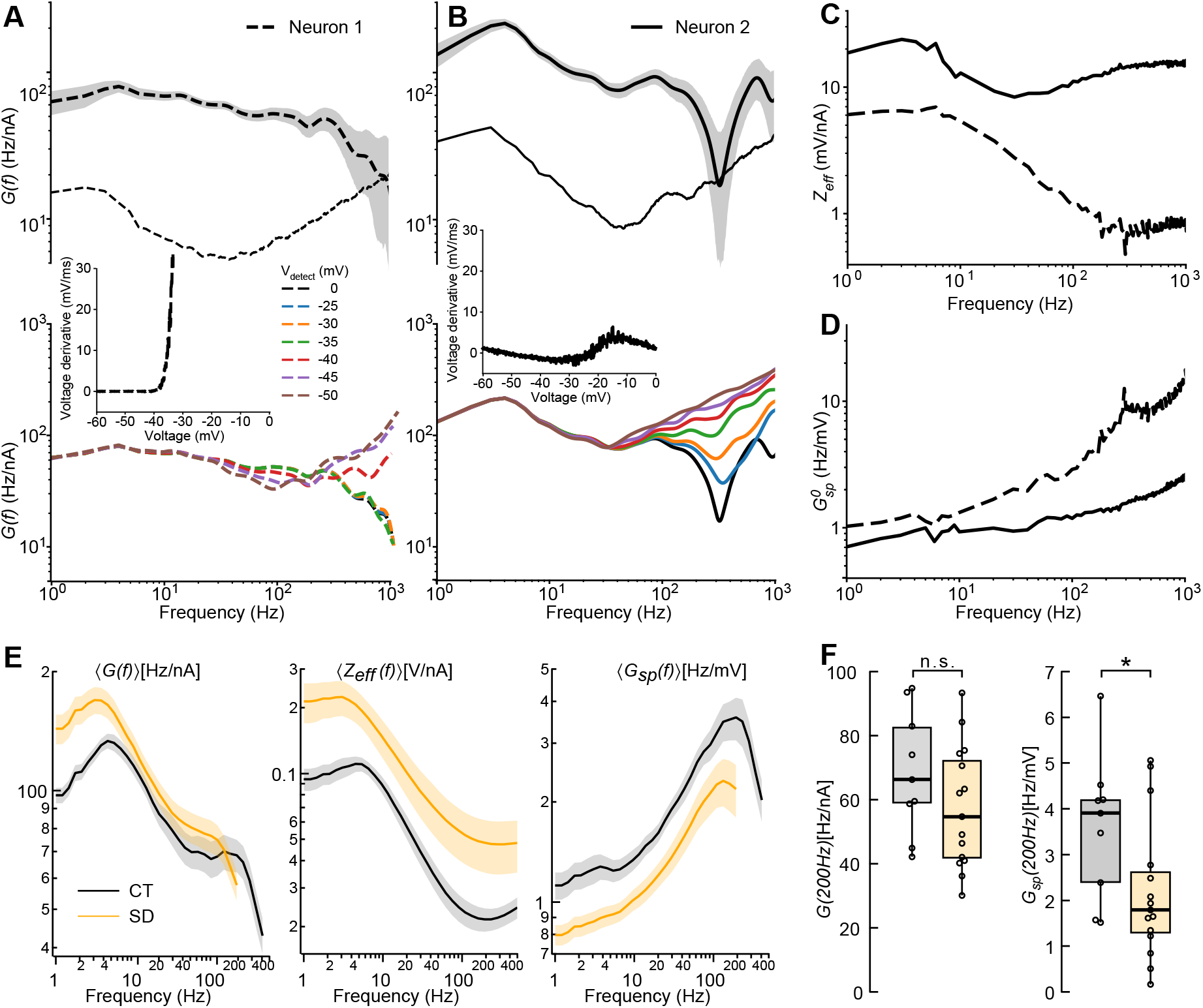
Ischemic insult slows AP initiation. **A** Dynamic gain function *G*(*f* ) (upper panel) of one pyramidal neuron from the treatment group in Revah et al. (2019), and the associated gain curves recalculated with lower AP detection thresholds ranging from 0 mV to -50 mV (lower panel). Inset panels are the phase plots estimated by averaging voltage derivatives at each voltage value. Note the wide bandwidth of the original gain function. Grey areas are the 95th percentile bootstrapping confidence intervals. Thin line represents the 95th percentile noise floor (see Materials and Methods). **B** as A, but for another neuron from the treatment group with drastically different gain. Firing rates are 4.10 Hz and 4.68 Hz in A and B respectively. **C** and **D** Effective impedances and zero-delay spike gains of the dynamic gain functions in A and B. 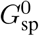 is the ratio between zero delay dynamic gain (*V*_detect_ = -50 mV in C and D) and effective impedance. **E** Grand average of *G*(*f* ), *Z*_eff_, and *G*_sp_(*f* ) for control neurons (CT, n=9) and neurons undergoing brief hypoxia and spreading depolarization (SD, n=15). Lines and shades represent means and their standard error. *G*(*f* ) and *G*_sp_(*f* ) are displayed for the frequency region in which all individual traces are significant, i.e. above the respective noise floor. **F** *G*(*f* ) at 200 Hz does not differ between groups. Due to the large *Z*_eff_ differences between the groups, the spike gains at 200Hz are significantly different (t-test, *p=0.048).

As a first step, we estimated the phase plot from the recordings (see Materials and Methods) by averaging voltage derivatives at different voltages (inset panels in A and B). This reveals a difference in threshold voltage, but most importantly, a large difference in the amount of voltage-sensitive depolarizing currents between the two neurons. For neuron 1, the voltage derivative changed little for voltages below -40 mV, then suddenly increased to 30 mV/ms within 8 mV, reflecting large AP initiation currents. Neuron 2, in contrast, had smaller depolarizing currents and a higher threshold. The position of the local minimum *V*_loc_ was around -30 mV for neuron 2 and, although it could not be discerned precisely, it was likely just below -40 mV for neuron 1.

In the second step, we reduced the AP initiation delay by lowering the AP detection threshold *V*_detect_ analogous to the analysis of Eyal’s model in Fig. 4B. The recalculated dynamic gain functions for the various *V*_detect_ values are given in the lower panels of Fig. 8A and B. For neuron 1, the gain curves changed abruptly, once *V*_detect_ reached -40 mV and already at -45 mV a saturation was reached. Apparently, only this narrow supra-threshold voltage range contributed to the uncertainty in AP timing. This corresponds to the steep change in the phase plot (inset). The initiation-related decay of the dynamic gain affected only frequency components above 300-350 Hz. When the AP was detected around *V*_loc_, the dynamic gain curve resembled that of a LIF neuron model without voltage-dependent initiation and with a hard threshold Brunel et al. (2001). This result replicates the simulations shown in Fig. 4B, albeit with an almost four times higher bandwidth of the dynamic gain decay, as compared to Eyal’s type II model.

In neuron 2, the recalculated dynamic gain curves behaved differently. The transition to LIF-like behavior was much more gradual (Fig. 8B, colored traces). The uncertainty in AP timing accumulated during the transition through a much larger voltage range from above -25 mV down to -40 mV, mirroring the more gradual change in phase plot slope (inset). Another important difference is the affected frequency range. In neuron 2, suprathreshold delay variability already reduced the encoding for frequencies above 90-100 Hz, similar to what we observed for Eyal’s model.

Our insights into the different suprathreshold dynamics are complemented by an analysis of the subthreshold contribution to dynamic gain. We calculated the effective impedance as the ratio of Fourier transform of voltage and Fourier transform of current (see Materials and Methods). The two neurons’ effective impedances and zero-delay spike gains are juxtaposed in Fig. 8C and D. This reveals the considerably larger effective impedance of neuron 2 as the origin of this neuron’s large absolute dynamic gain value. The fact that this effective impedance rises for frequencies above 30 Hz is puzzling and we can only speculate about a contribution of sodium channels, possibly due to the increased threshold in neuron 2. For neuron 1, the effective impedance approximates the shape one would expect from a passive soma with dendrites. It is dominated by a cutoff of 12.8 Hz, i.e. a membrane time constant of 12.5 ms. Beyond the cutoff, the slope is less negative than -1, most likely due to the presence of dendrites. The zero-delay spike gains of the two neurons are rather similar at low frequencies, but above 20 Hz, they deviate increasingly. Because these curves are obtained by dividing zero-delay dynamic gain curves by the effective impedance, this deviation originates in the unusual impedance increase in neuron 2.

In summary, the differences between the two neurons’ dynamic gain curves can be attributed to subthreshold and suprathreshold contributions as follows: the difference in absolute values is caused by the higher effective impedance of neuron 2. The different locations of the dynamic gain drop, and consequently the different bandwidths are explained by the substantially slower AP initiation in neuron 2. It causes an earlier drop in dynamic gain due to larger initiation delay variability. The remaining difference is the striking rise in the high-frequency region of the second neuron’s dynamic gain. We suspect that at the high frequencies the extrinsic, stimulation-induced currents dominate the second neuron’s weaker intrinsic currents. Consequently, the exact time of threshold crossing is influenced by high-frequency input fluctuations, similar to the situation in a LIF neuron model. This is the reason for the rising dynamic gain and the relatively narrow confidence interval. In the context of the original study Revah et al. (2019), it is interesting to note, that cells with such a weak intrinsic AP initiation appeared only after hypoxia and spreading depolarization, which also compromised the molecular integrity of the axon initial segment.

Applying dynamic gain decomposition to all the cells in the original data set, we could further investigate the previously described effect of hypoxia-induced spreading depolarization on the dynamic gain. The grand averages for dynamic gain ⟨*G*(*f* )⟩, effective impedance ⟨*Z*_eff_(*f*) ⟩ and spike gain ⟨*G*_sp_(*f*) )⟩ were calculated over the frequency range, where data where available from all cells. For ⟨*G*(*f*)⟩ and *G*_sp_(*f*) that only includes the range up to 300 Hz, at which point at least one cell’s *G*(*f*) touched the noise floor. The grand averages, together with their standard errors are shown in Fig. 8E. They show clearly that the average effective impedance is increased in the treated cells, and that this difference dominates the unnormalized dynamic gain curves. While the dynamic gain of the treated cells appears to drop earlier, between 100 and 200 Hz as compared to the control cells >300 Hz, this difference does not manifest in a statistically significant difference. The dynamic gain values at 200 Hz are spread out over similar ranges for both groups (Fig. 8F).

In contrast, the impedance-corrected spike gain values are significantly different (Student’s t-test, p=0.048). These examples show that all aspects of dynamic gain decomposition can be applied to experimental data and serve to suggest biophysical parameters as the basis of individual dynamic gain features.

## Discussion

Here, we developed a straightforward method to decompose the contribution of subthreshold and suprathreshold dynamics on population encoding. It allows for a unified and comprehensive approach of dynamic gain analysis because it can be applied to simulated data from models of any complexity and even measurements from real neurons. When we applied the dynamic gain decomposition to a complex multi-compartment model equipped with a biophysically plausible AP initiation mechanism, we found that the model’s high encoding bandwidth is mainly enabled by a high potassium conductance around threshold. This also causes a type II excitability, but the tendency to high-frequency bursting is not in itself beneficial for a high bandwidth. The decomposition also guides the definition and choice of working points at which model variants or cell types can be compared without bias. Iso-(voltage-)fluctuation and iso-(initiation-)delay working points control for sub- or suprathreshold contributions, respectively. We found the iso-delay criterion particularly useful for understanding how changes in input correlation and dendrite size affect encoding bandwidth. We also applied the decomposition to recordings from two layer 5 pyramidal neurons, which were differently affected by hypoxia. Their different dynamic gain shapes and bandwidths resulted from striking differences in effective impedance and intrinsic initiation currents. This demonstrates that dynamic gain decomposition can connect cellular physiology to network function.

### Dynamic gain function components

Two of the three components we introduced (effective impedance and dynamic gain decay) capture well-defined subthreshold and suprathreshold processing steps and are independently calculated. The third component (zero delay spike gain) represents dynamic gain features not captured by the other two. Effective impedance and zero-delay spike gain jointly capture the impact of subthreshold dynamics on population encoding. Their product represents a LIF-like dynamic gain function. The dynamic gain decay in the high frequency region, reflects how the AP initiation dynamics limits the stochastic input’s control over the precise AP time. The identification of separate components underlies the idea to align working points across model variants before dynamic gain decomposition. A related approach, a dynamic gain decomposition into separate components, lies behind the approach to classify APs into few groups, and calculate the group-based dynamic gain components. Here, this approach elucidated the individual contributions of APs leading to a burst, earlier we applied it to isolate the effect of AP timing relative to slow input oscillations Merino et al. (2021).

### Fixing working points isolates parameter effects

The dynamic gain function is not a fixed curve but depends on the working point of the neuron model. It can change substantially when the operating regime is changed from mean-driven to fluctuation-driven (Fig. 3G). Thus, it is essential to properly align the working points of model variants, such that the differences detected in the dynamic gain functions originate mainly from the targeted parameter change, rather than unrelated working point factors. One criterion we adopted here is fixing the firing rate at 5 Hz. A second criterion can be chosen to fully determine the working point on the two-dimensional manifold of firing rate, defined by *μ*_*I*_ and *σ*_*I*_ (Fig. 3D). Previous studies have focused on fixing the subthreshold dynamics, either by fixing the voltage fluctuations *σ*_*V*_ Tchumatchenko et al. (2011); Köndgen et al. (2008); Ilin et al. (2013), or the firing pattern (*CV*_*ISI*_) Vilela and Lindner (2009); Lazarov et al. (2018); Zhang et al. (2022). Both criteria, at least in part, relate to the subthreshold voltage fluctuation. Keeping them constant is suitable if the major differences between the compared models manifest mostly in the suprathreshold regime. Here, we introduced a complementary criterion, the initiation delay. By fixing the mean of this random variable, we managed to control the impact of the suprathreshold AP dynamics across models, which corresponds to the dynamic gain decay in the high frequency region. We found this criterion particularly useful for studying the Brunel effect and the dendritic effect on Eyal’s model, since both mainly affect the subthreshold dynamics. What is more, we generalized the approach to LIF-like models, by fixing the average voltage derivative at threshold (see the Extended Data Fig. 5 F).

### Uncovering the role of AP initiation dynamics for high-frequency dynamic gain

Lowering the AP detection voltage to the voltage range where intrinsic currents and external input jointly govern the voltage dynamics, results in changes to the dynamic gain in the high-frequency region until it resembles that of a LIF-like neuron. Obviously, this AP detection at low voltages does not represent a physiological mechanism; an AP is special precisely because its strong depolarization does gate other ion channels. However, the virtual AP times resulting from the low detection threshold still reveal a physiologically meaningful property. Because dynamic gain measures the susceptibility of AP times to extrinsic perturbation, the difference between dynamic gain curves obtained with high and low detection thresholds uncovers how this susceptibility is built up or eroded during the AP initiation interval. This simple, phenomenological approach reveals how the AP initiation dynamics within different voltage regions contributes to the physiologically meaningful dynamic gain. Such information is otherwise inaccessible, except for the most simple intrinsic dynamics, when it can be analytically calculated Fourcaud-Trocmé et al. (2003); Wei and Wolf (2011). In models and experimental data, we observed the same, expected relation between the initiation-related dynamic gain modulation and the voltage dependence of initiation currents. The steeper d*V/*d*t* rises, the larger is the bandwidth and the smaller is the range of detection thresholds for which the dynamic gain shape changes (see Fig. 4A and 4E; Fig. 8A and 8B). Our analysis of Eyal’s model revealed that the working point strongly influences this initiation-related dynamic gain modulation. However, the model’s encoding bandwidth is never chiefly limited by the initiation dynamics, because the subthreshold signal transformations, the effective impedance *Z*_eff_ and zero-delay spike gain 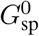, impose a cut-off at lower frequencies. This is in contrast to the findings in cortical pyramidal cells, where the initiation dynamics seems to pose the ultimate limitation for the encoding bandwidth (Fig. 8A).

This analysis approach holds great promise for future studies for two reasons. First, it provides a common basis to disentangle the role of initiation dynamics in all models and experimental data. Second, it is sensitive to the input statistics. It changes when the neuron is studied at a different working point and thereby the relation between extrinsic and intrinsic currents shifts. In that respect, the dynamic gain decay analysis is much more informative than the conventional metrics of initiation dynamics, the AP phase plot.

### Implications for neuronal network modelling

The shape and bandwidth of the dynamic gain play important roles for studies of information processing in large, recurrent neuronal networks. In large network simulations, the neuron models should therefore feature realistic dynamic gain functions with realistic dependencies on the input statistics. However, on the scale of tens of thousands of neurons, the computational costs of multi-compartment models are almost prohibitive. Our dynamic gain decomposition suggests routes to create simpler models with realistic dynamic gain curves that depend on input statistics just as the full multi-compartment models do. Point neuron models can be combined with analytical input transformations, similar to the work of Aspart and colleagues Aspart et al. (2016). The realistic, limited bandwidth could be obtained by adding a random AP initiation delay distribution. The subthreshold influence on the dynamic gain curve could be created by a few voltage-gated or AP time-dependent conductances that create adaptation and resonances. The electrotonic separation between soma and AIS, which shapes the signal arriving at the AP initiation zone Brette (2013); Zhang et al. (2022), could be represented in three-compartment models, representing AIS, soma, and dendrite, similar to the two-compartment model utilized by Ostojic and colleagues Ostojic et al. (2015). Those approaches could provide realistic dynamic gain curves at lower computational cost.

### Cellular physiology of dynamic population coding

Because dynamic gain decomposition works along the biophysical signal transformation cascade, its results can be interpreted in physiological terms and corroborated with conventional electrophysiological methods, such as potassium channel pharmacology Higgs and Spain (2011). This is particularly helpful in the emerging fields of axon initial segment plasticity and spectrinophaties, where structural changes at the site of AP initiation are observed as consequences of altered input Grubb and Burrone (2010); Grubb et al. (2011); Kuba et al. (2010); Jamann et al. (2021) or mutated structural molecules Parkinson et al. (2001); Lazarov et al. (2018); Wang et al. (2018), and the consequences for excitability and encoding capacity need to be quantified in order to understand the functional impact on the population level. We propose that future studies of axon initial segment plasticity can use dynamic gain measurements at carefully maintained working points to quantify potential circuit-level consequences. Dynamic gain decomposition can then be used to inform mechanistic models of how the observed molecular and structural rearrangements affect functional differences or, alternatively, maintain encoding precision while changing excitability. Foundations for such a connection between physiology and encoding precision have already been laid by studies that tie structural changes at the axon initial segment Lazarov et al. (2018); Revah et al. (2019) or dendrite size Testa-Silva et al. (2014); Goriounova et al. (2018); Ostojic et al. (2015) to changes in the dynamic gain. It will be possible to apply the decomposition method to these recently obtained data, to attribute the observed changes to sub- and suprathreshold signal transformations, and thereby critically test the concepts formulated in those studies. The decomposition examples in this study reveal that unlike Eyal’s model, the real neuron’s encoding bandwidth is limited by the initiation dynamics. These preliminary results opens up new questions for future research, because the biophysical origin of the effective impedance’s peculiar shape is unclear, as is the wide bandwidth of the initiation-related dynamic gain decay (Fig. 8A-C). Ultimately, to understand population encoding in the brain, we need a better understanding of the physiological working points, experienced by neurons *in vivo*.

## Acknowledgements

This work was supported by the Ministry for Science and Culture of Lower Saxony (MWK) and VolkswagenStiftung through the program “Niedersächsiches Vorab”. We acknowledge further support through the China Scholarship Council (to CZ), the Bundesministerium für Bildung und Forschung (BMBF, Federal Ministry of Education and Research) grant 01GQ1005B (FW, AN), the VolkswagenStiftung under grant no. ZN2632 (to FW), and through the Deutsche Forschungsgemeinschaft (DFG, German Research Foundation) through CRC 1286 and SPP 2205 (FW) and through grant 436260547 (AN, FW), in relation to NeuroNex (NSF 2015276). The funders had no role in study design, data collection and analysis, decision to publish, or preparation of the manuscript.

## Extended data

**Figure 3-1:**
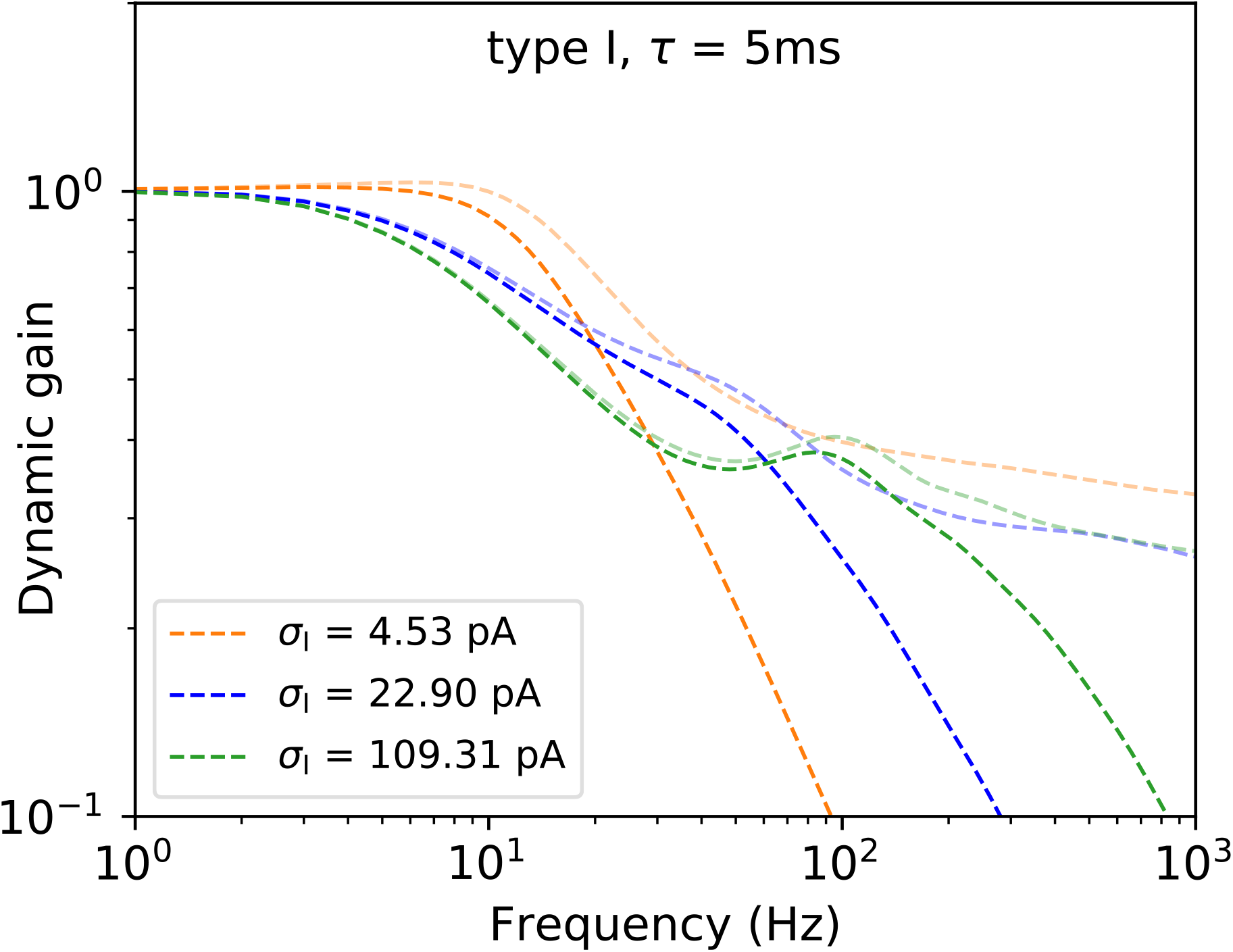
Dynamic gain functions and corresponding zero-delay dynamic gain functions of Eyal’s type I model at three working points ranging from nearly mean-driven to very fluctuation-driven. The three working points are those provided in the main text for figure 3. Slow AP initiation dynamics limits the encoding bandwidth only when the working point is mean-driven (orange). In the the fluctuation driven cases, substantial deviations between dynamic gain and zero-delay gain appear only at frequencies above the cut-off frequency.

**Figure 6-1:**
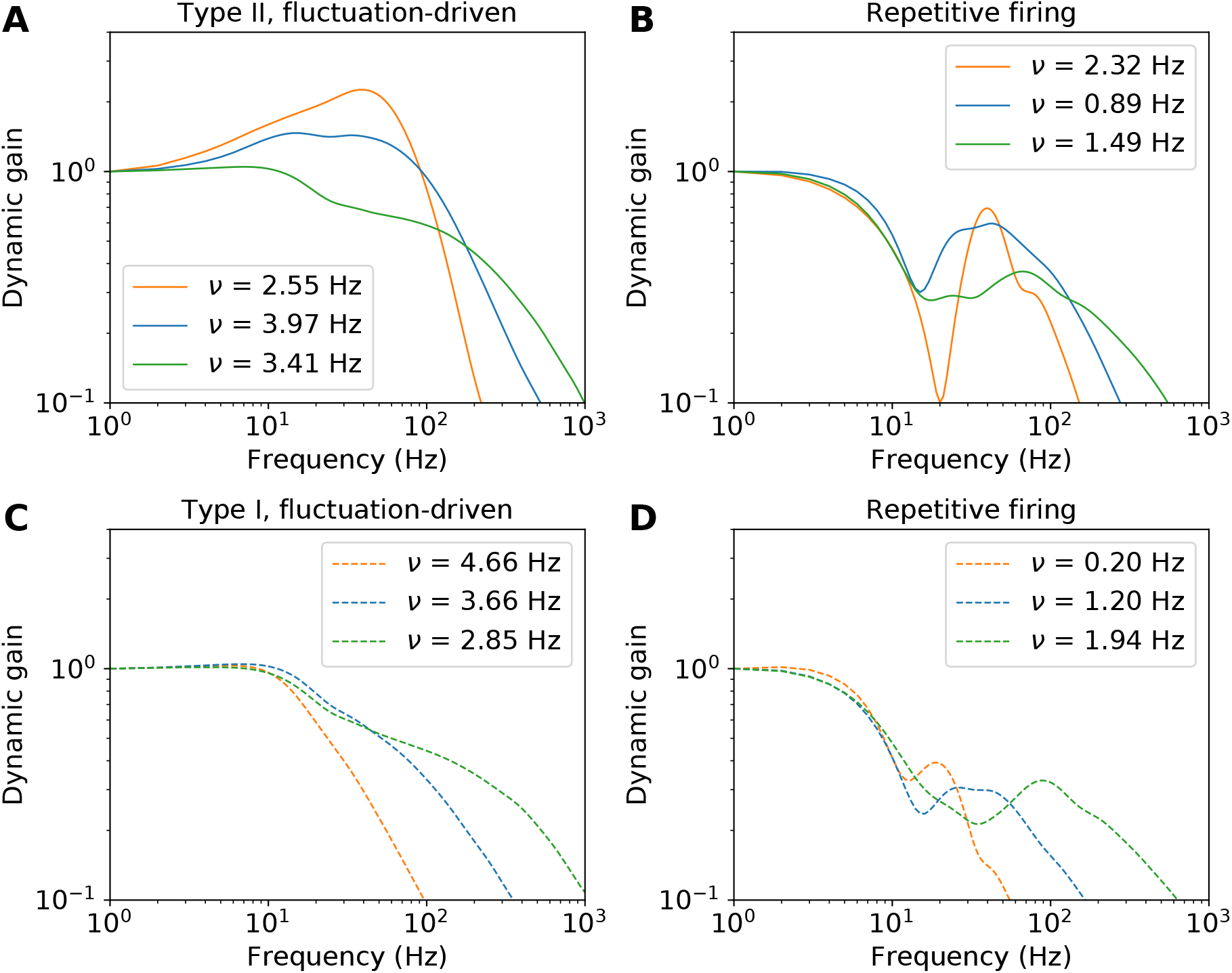
Dynamic gain functions of fluctuation-driven APs and repetitive firing APs of type II and type I models, at the three working points ranging from mean-driven to very fluctuation-driven. *τ* = 5 ms.

**Figure 7-1:**
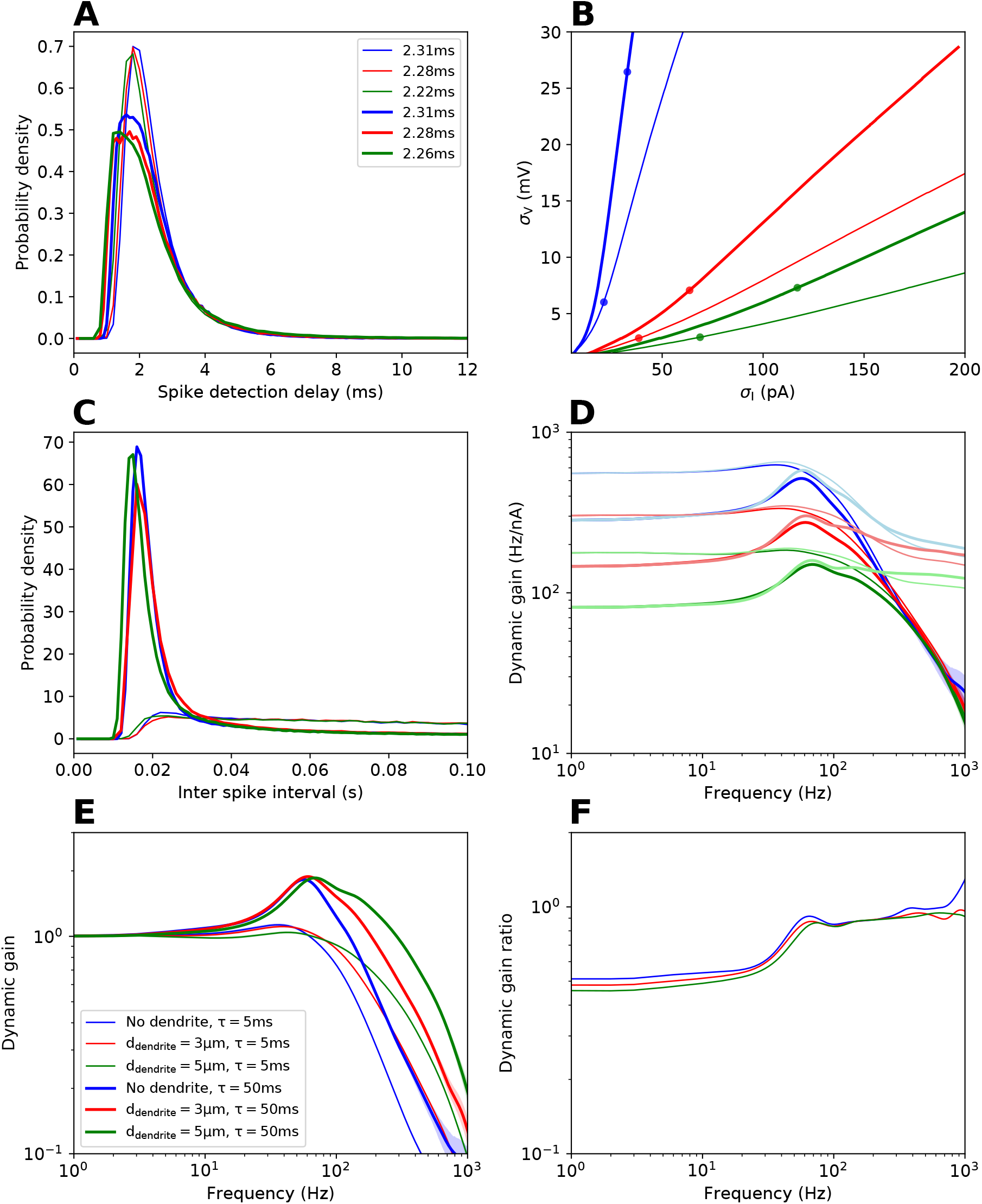
Dynamic gain functions of Eyal’s type II models with different dendrite sizes at iso-delay working points. This figure complements the data, shown in figure 7 by adding responses for *τ* = 50 ms (wide lines) to the 5 ms data (thin lines). In both cases, a larger dendrite can enhance high frequency encoding, while the Brunel effects are almost identical for the three model variants.

